# In depth analysis of kinase cross screening data to identify chemical starting points for inhibition of the nek family of kinases

**DOI:** 10.1101/137968

**Authors:** Carrow I. Wells, Nirav R. Kapadia, Rafael M. Couñago, David H. Drewry

**Affiliations:** Structural Genomics Consortium, UNC Eshelman School of Pharmacy, University of North Carolina at Chapel Hill, NC 27599 USA.; Structural Genomics Consortium, Universidade Estadual de Campinas – UNICAMP, Campinas, SP, 13083 Brazil.

## Abstract

Potent, selective, and cell active small molecule kinase inhibitors are useful tools to help unravel the complexities of kinase signaling. As the biological functions of individual kinases become better understood, they can become targets of drug discovery efforts. The small molecules used to shed light on function can also then serve as chemical starting points in these drug discovery efforts. The Nek family of kinases has received very little attention, as judged by number of citations in PubMed, yet they appear to play many key roles and have been implicated in disease. Here we present our work to identify high quality chemical starting points that have emerged due to the increased incidence of broad kinome screening. We anticipate that this analysis will allow the community to progress towards the generation of chemical probes and eventually drugs that target members of the Nek family.

## Introduction

Protein kinases are a group of enzymes that catalyze the transfer of a phosphate group from ATP to serine, threonine, and tyrosine residues of substrate proteins. This simple reaction modifies the shapes and activities of target proteins and is a key mechanism for signal transduction. Accordingly protein kinases are implicated in a multitude of pathophysiological conditions and are attractive drug targets.^1^ There are approximately 518 protein kinases in humans and a small percentage of them have been the subject of extensive research by pharmaceutical and academic scientists. Kinases are tractable drug targets, and 28 small molecule kinase inhibitors are approved for therapeutic use.^2^ Unfortunately the majority of the research has enlightened the function of only 20% of the kinases, leaving the biology of the remaining 80% relatively dark and poorly understood.^3^

Our group is currently developing a comprehensive kinase chemogenomic set (KCGS) to aid in the elucidation of the biological function and therapeutic potential of human protein kinases.^4^ To accomplish this goal we are building a set of potent, narrow spectrum kinase inhibitors that together cover the whole kinome. One strategy to develop these high quality inhibitors is to work on families of these understudied dark kinases. One such family that has experienced very little targeted medicinal chemistry effort is the NIMA-related Kinase (Nek) family of serine/threonine kinases. Table 1 lists the number of citations in PubMed as a simple metric to rank the amount of research interest a kinase target has received. Nek2 has received the most attention in the Nek family, Nek5 has garnered the least interest, and all are understudied. The number of citations for ERBB2^5^, a well studied and highly cited kinase, is included in the table to provide some context. The founding member of the Nek kinases is the Never In Mitosis A (NIMA) protein of *Aspergillus nidulans*. NIMA plays a critical role in mitosis in *A. nidulans*, wherein degradation of NIMA is essential for mitotic exit. In line with its critical role, NIMA-related kinases have been found in higher organisms such as *plasmodium*, *drosophila*, *xenopus*, mice and humans. In humans, the Nek family is comprised of 11 members – Nek1 to Nek11. As observed with NIMA, Nek family members are involved in various stages of cell cycle regulation.^6–9^ Given that they play roles in cell cycle regulation, these kinases have the potential to become important therapeutic targets for a range of diseases (Table 1). To realize this potential, a greater understanding of the roles each Nek plays in physiology and pathophysiology must be elucidated. Here we briefly describe some of the known biology of the Nek family members, and then identify potential chemical starting points for synthesis of narrow spectrum small molecule inhibitors. Though very little chemistry effort has been expended on the Nek family (Nek2 being the only exception), careful analysis of published kinase cross screening data from projects focused on other kinases allows us to highlight nascent structure activity relationships (SARs) that point to viable Nek inhibitor starting points for focused medicinal chemistry efforts.

**Table 1.**
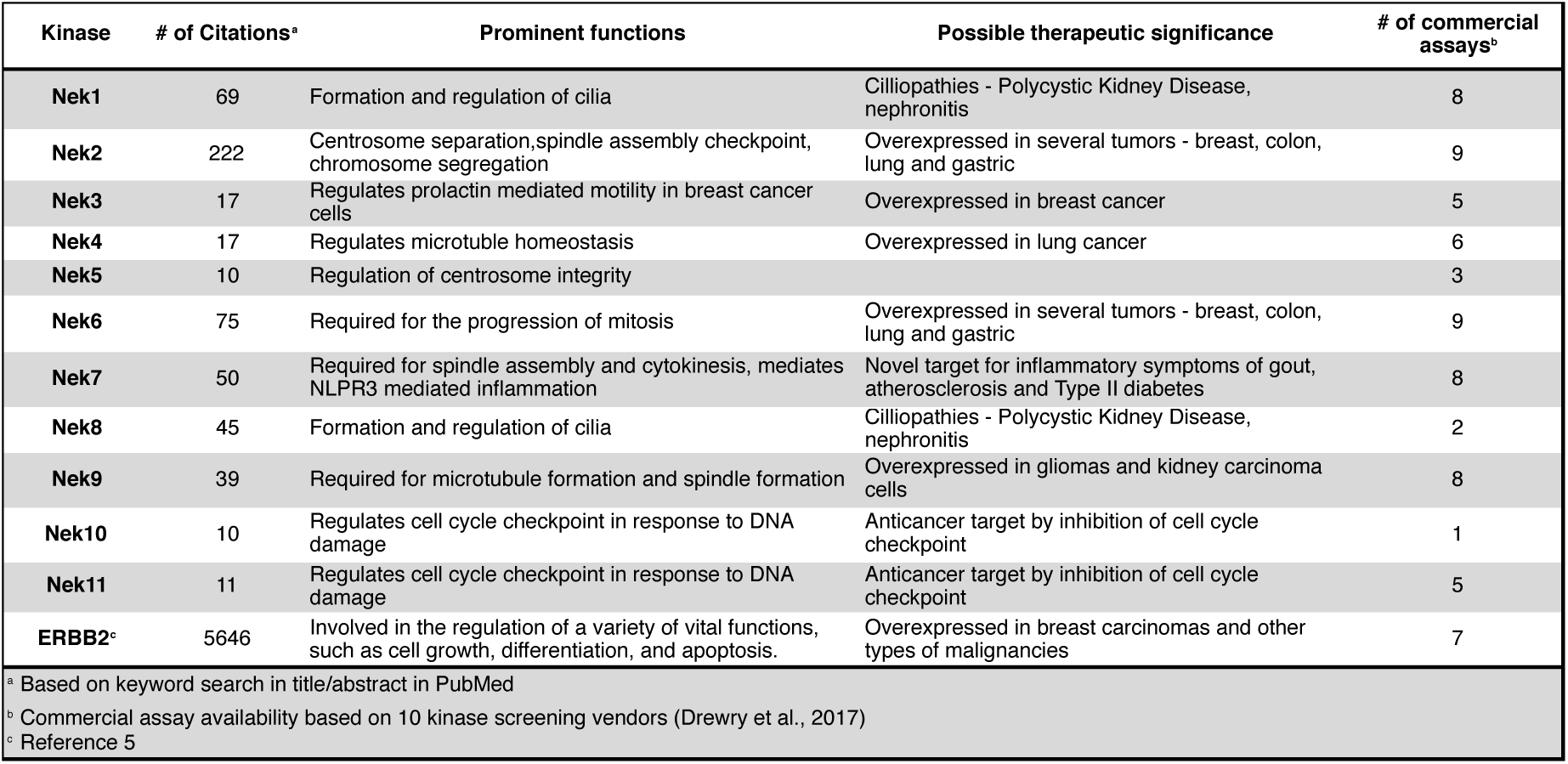
Biological function, disease relevance, and assay availability for members of the Nek kinase family.

### Nek family Active Site Residue Comparisons

To assess whether selectivity between Nek family members would be problematic we compared 28 kinase active site residues^10^ from all eleven Nek family members in a pairwise manner (Figure 1a; Table S1). From this analysis it was readily apparent that Nek10 is the most divergent from the other Nek family kinases as indicated by both the yellow horizontal and vertical line. Nek6 and Nek7 show the highest degree of sequence identity with only one of the 28 active site residues we investigated differentiating them from each other. This analysis suggests that obtaining inhibitor selectivity between Nek 6 and Nek7 could prove to be challenging. To get a feel for this based on actual inhibition data, we calculated the activity homology between each pair of Nek kinases.^11^ Activity homology is a measure of the likelihood that a kinase inhibitor that inhibits a particular kinase will also inhibit another kinase. We have chosen 80% inhibition as our cutoff here. We identify all compounds that inhibit a particular Nek ≥80%, and then calculate what percentage of those compounds inhibit each other Nek ≥80%. This activity homology calculation allows quick visualization of where there is inhibitor selectivity between Nek family kinases (Figure 1). We find the prediction based on close similarity of active site residue that Nek 6 and Nek 7 selectivity may be difficult to obtain to be true when looking at activity homology between Nek6 and Nek7 using recently published broad screening results from the Published Kinase Inhibitor Set 2 (PKIS2) (Figure 1b). 88% of the compounds that inhibited Nek6 at 80% inhibition or greater at 1 µM also inhibited Nek7 at 80% inhibition or greater. Nek5 and Nek1 are also fairly similar by active site sequence analysis (24/28 residues in common) implying that selectivity could be difficult to achieve. However, comparison of activity data from PKIS2 compound screening revealed that 29 compounds showed ≥80% inhibition for Nek5, and only 5 of these compounds showed ≥80% inhibition on Nek1. Several of these five compounds, though, have very broad kinase inhibition profiles. Our analysis showed that although Nek4 has 60-70% active site sequence similarity with Nek1, Nek2, Nek3, and Nek5, it has very low hit homology with these other members of the Nek family. However, this result may be confounded by the fact that Nek4 has a very low hit rate in the PKIS2 dataset (Table 2) with only 5 compounds that showed greater than 80% inhibition on Nek4.

**Figure 1.**
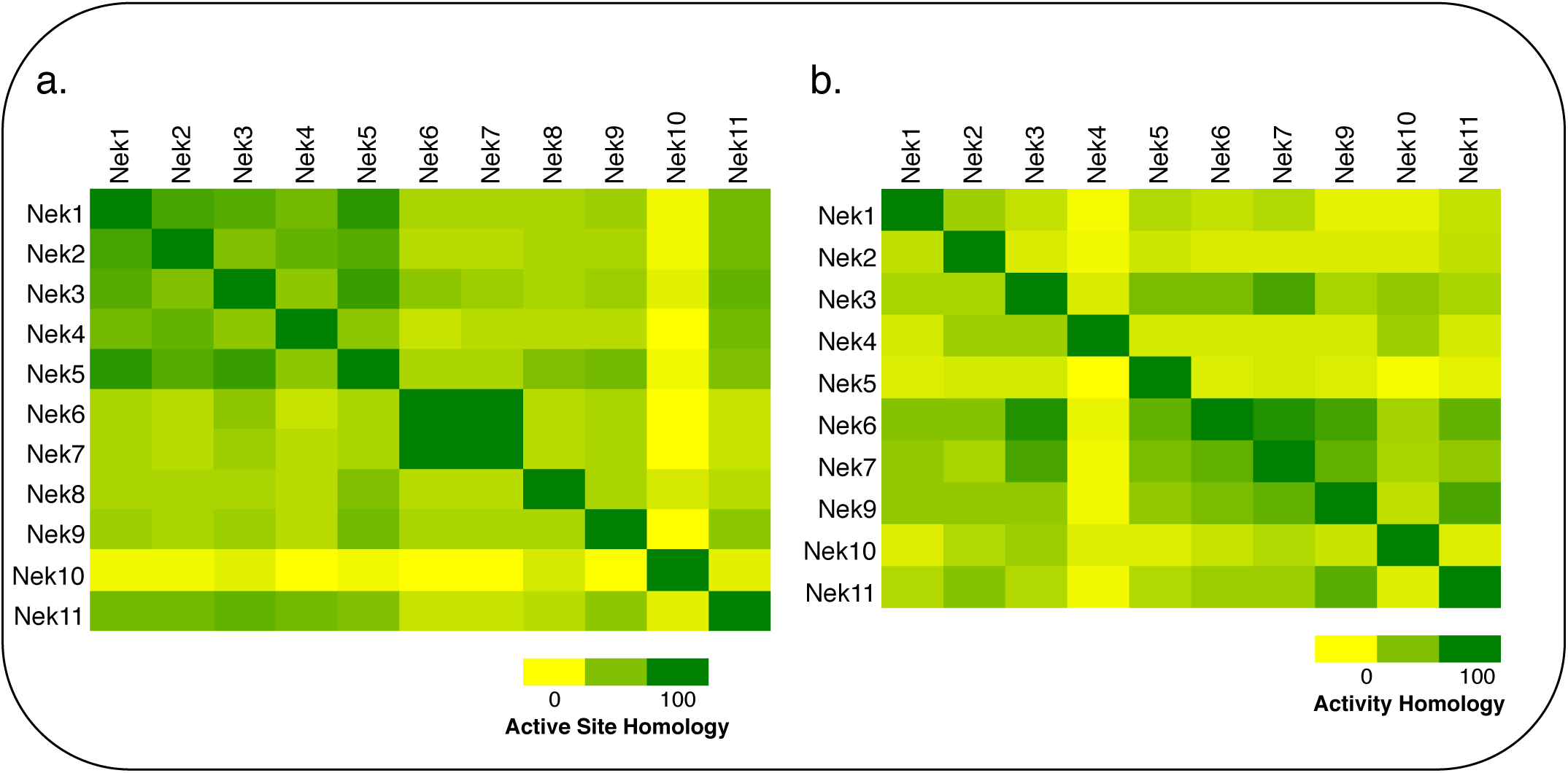
Pairwise comparisons of the **a**. active site sequence identity **b**. activity homology calculated using data from PKIS2^4^

**Table 2.** Nek family kinase selectivity. Selectivity scores for the Nek family based upon screening data from large published screening sets with FLT3 value for comparison.

| Compound set | PKIS<sup>a</sup> | PKIS2<sup>b</sup> | BMS<sup>c</sup> | RBC<sup>d</sup> | Ambit<sup>e</sup> | GSK<sup>f</sup> | Millipore<sup>g</sup> |
| --- | --- | --- | --- | --- | --- | --- | --- |
| no. of cmpds | 368 | 645 | 21851 | 178 | 72 | 577 | 158 |
| Nek1 | 0.011 | 0.051 | 0.001 | 0.051 | 0.014 | – | – |
| Nek2 | 0.005 | 0.057 | 0.003 | 0.006 | 0.056 | 0.059 | 0.006 |
| Nek3 | – | 0.031 | – | 0.006 | 0.014 | – | 0 |
| Nek4 | – | 0.012 | – | 0.039 | 0.014 | – | – |
| Nek5 | – | 0.065 | 0.0286 | – | 0.014 | – | – |
| Nek6 | 0 | 0.023 | 0.017 | 0 | 0.042 | 0.031 | 0 |
| Nek7 | 0 | 0.035 | 0.0315 | 0 | 0.042 | 0.035 | 0 |
| Nek8 | – | – | – | – | – | – | – |
| Nek9 | 0.033 | 0.051 | 0.009 | 0.028 | 0.042 | 0.016 | – |
| Nek10 | – | 0.059 | – | – | – | – | – |
| Nek11 | – | 0.031 | – | 0.011 | 0.042 | – | 0.095 |
| FLT3 | 0.14 | 0.23 | 0.15 | 0.41 | 0.036 | 0.14 | 0.3 |

<sup>a</sup>values for PKIS calculated by the number of compounds that inhibited each Nek by 50%≤ divided by the total number of compounds screened (Elkins et al., 2016)
<sup>b</sup>values for PKIS2 calculated by the number of compounds that inhibited each Nek 50%≤ divided by the total number of compounds screened (Drewry et al., 2017)
<sup>c</sup>values from Bristol-Myers Squibb screening data (hit rate calculated based on <13% of control) (Posy et al., 2011)
<sup>d</sup>values from Reaction Biology Corporation data (Anastassiadis et al., 2011)
<sup>e</sup>values calculated from reported Ambit Kd values - number of compounds that inhibit each Nek ≥500nM divided by the total number of compounds (Davis et al., 2011)
<sup>f</sup>values from GlaxoSmithKline screening taken from Nc≤1 (Bamborough et al., 2008)
<sup>g</sup>values calculated using EMD Millipore screening data. Number of compounds that inhibit each Nek 50%≤ divided by total number of compounds screened (Gao et al., 2013)

### Chemical Tractability of the Nek family

Since we are interested in identifying high quality chemical starting points for the Nek family of kinases, and there are no medicinal chemistry papers published on any member of the Nek family other than Nek2, we analyzed publically accessible data from large screening sets that have been described in the literature.^4^, ^11–16^ A number of groups from large pharmaceutical companies, biotech companies, screening vendors, and academic labs have published results from screening sets of kinase inhibitors against broad panels of kinase assays. These studies differ in size and chemical diversity of the small molecule screening set, concentration of inhibitor utilized for screening, type of data collected (single shot %I or K_d_/IC_50_), format of the kinase assays (binding versus enzyme inhibition), and number of kinases included in the exercise. In spite of these differences, common threads emerge. The most important outcome for our purpose is that compounds made for one kinase often inhibit other kinases, and this information, when shared like this, can be exploited to identify chemical matter for kinases that have received scant formal attention. We compiled this information and examined the hit rate for each Nek family member screened in this manner so that we could get a feel for the overall chemical tractability of the Nek family. How hard is it to find small molecule inhibitors? The BMS publication reports on the largest number of compounds, almost 22,000, but only six out of the eleven Nek family kinases were screened. The Nek kinases that were screened had relatively low hit rates, typically about 3% or less with the majority being under 1% (Table 2). The large size of the compound set likely gives a fair indication of tractability with current chemical matter. We have included hit rates for the kinase FLT3 in the table for comparison. FLT3 is a kinase that is usually among the most easily inhibited in the published broad compound profiling efforts. We also evaluated the Nek family hit rate from the PKIS2 screening set and saw that there were significantly higher hit rates on Nek5 than in the other publically available data sets (Table 2, Figure S2). This observation is an exciting finding because there are no published selective and potent Nek5 inhibitors and it could provide chemical starting points for the synthesis of inhibitors for inclusion into the KCGS that we are building. Nek8 has not been profiled in any public report, so no hit rates are reported. This dark kinase is one where the identification of any inhibitor would be impactful. It is also of note that even though this data was compiled using assay formats from four different vendors (Nanosyn, DiscoverX, RBC, and Millipore) and a variety of compounds and compound panel sizes, the hit rates are relatively consistent across the respective Nek kinases. This demonstrated that hits can be found for all Nek family members that have been screened, but overall the Nek family members have lower hit rates than the average kinase.

### Nek Inhibition by Promiscuous Kinases Inhibitors

When looking for starting points for Nek inhibitors, a subset of molecules we identified were promiscuous kinase inhibitors. These promiscuous inhibitors have activity on a large percentage of the kinome including the Nek family. This observation provides confidence that the Nek family of kinases can be inhibited with known chemical matter. Even if the compounds are not selective enough to be used as tools, these results identify features that confer activity. Tamatinib (R406), originally published as a SYK inhibitor,^17^ inhibits 8 of the 9 Nek kinases it has been evaluated on below 500 nM. It does, however, inhibit 34% of the kinases it has been screened against with Kd values below 300 nM.

**Table 3.** Promiscuous kinase inhibitors that inhibit Nek kinases

| Compound | Tamatinib<sup>a</sup> | KW-2449<sup>a</sup> | AST-487<sup>a</sup> | EXEL-2880<sup>a</sup> | TG-101348<sup>a</sup> | PD-173955<sup>a</sup> | cmpd 2516<sup>b</sup> | cmpd 3177<sup>b</sup> | GSK1059615<sup>c</sup> |
| --- | --- | --- | --- | --- | --- | --- | --- | --- | --- |
|  | K<sub>d</sub> (nM) |  |  |  |  |  | IC<sub>50</sub> (nM) |  | % Control (10μM) |
| Nek1 | 170 | 3100 | 5200 | — | 3400 | 8300 | — | — | 47 |
| Nek2 | 260 | 5300 | — | — | 1700 | 110 | >10000 | 630 | 64 |
| Nek3 | 170 | 1000 | — | — | 640 | — | — | — | 77 |
| Nek4 | 1800 | 1100 | 460 | — | 1600 | — | 5 | 50 | 96 |
| Nek5 | 13 | 2300 | 1100 | — | 570 | 890 | — | — | 100 |
| Nek6 | 6.3 | 120 | 2500 | — | 120 | — | — | — | 81 |
| Nek7 | 11 | 210 | — | — | 160 | — | — | — | 88 |
| Nek8 | — | — | — | — | — | — | — | — | — |
| Nek9 | 16 | 520 | 1800 | 470 | 150 | — | — | — | 95 |
| Nek10 | — | — | — | — | — | — | — | — | 0 |
| Nek11 | 170 | — | — | — | 1200 | 3000 | — | — | 72 |
| Promiscuity | S(300nM)=0.34 | S(300nM)=0.26 | S(300nM)=0.26 | S(300nM)=0.28 | S(300nM)=0.18 | S(300nM)=0.21 | S(300nM)=0.26 | S(300nM)=0.36 | S(5%) at 10μM=0.17 |

<sup>a</sup> values from Davis et al., 2011 <sup>b</sup> values from Metz et al., 2010 <sup>c</sup> values from HMS LINC dataset ID 20085

There were also compounds such as KW-2449 that were broadly promiscuous but only potently inhibited one or two Nek family members. This compound was originally published as a FLT3 inhibitor,^18^ entered into Phase I clinical trials, and was later pulled from development^18^. This compound potently inhibits Nek6 and Nek7, which based on their sequence similarity would be expected to track together. Although the compound is broadly promiscuous it does not inhibit other Nek kinases, which indicates that selectivity between some family members can be achieved. There are also two promiscuous Type II kinase inhibitors, AST-487 and EXEL-2880, and each only inhibits one Nek family member, Nek4 and Nek9, respectively. This observation may be of interest since Type II inhibitors bind to the inactive or DFG-out conformation providing medicinal chemists additional options for building selectivity into these inhibitors.^19^ It has also been suggested that type II inhibitors in some cases may offer advantages over type I inhibitors, for example in residence time.

The screening that we performed on PKIS2 compounds also identified promiscuous inhibitors from several chemotypes that inhibited the Nek family of kinases (Figure 2). For example the 2,4-dianilinopyrimidine series is a known promiscuous chemotype that inhibits many kinases including every screened Nek family member under 10% of control at 1 µM. UNC5414, a compound from the pyrimidinyl pyrimidine series, also has a relatively broad inhibition profile but we do start to see some differentiation of Nek activity. Notably Nek4 was not inhibited, further supporting our conclusions from the Nek4 activity homology map (Figure 1b). Another interesting broad kinase inhibitor from PKIS2 is UNC5269. Although it inhibited almost half of the kinases tested, within the Nek family it only potently inhibited Nek4 and Nek10 (%control ≤10). These results, combined with docking models or crystallography, could provide insight into the preferred pharmacophores for inhibition preferred by specific Nek kinases.

**Figure 2.**
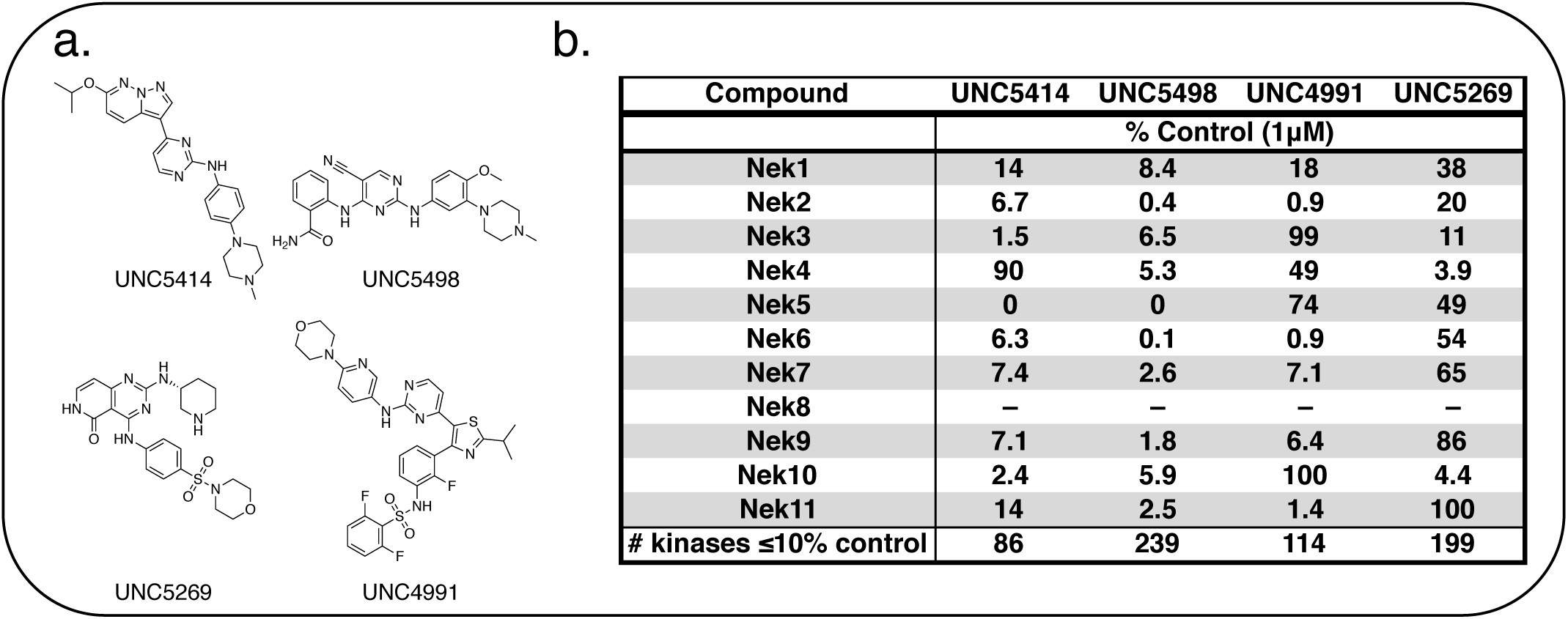
**a**. Structures of promiscuous kinase inhibitor exemplars from PKIS2 that target the Nek family **b**. Table of Nek family activity values as percent control

### Clinical compound targeting multiple Nek family members

We have identified one kinase inhibitor, Cyt387/Momelotinib (Figure 3), which is not broadly promiscuous, but still inhibited multiple members of the Nek family. Momelotinib was designed as a JAK2 inhibitor^20^ and has advanced into phase III clinical trials for myelofibrosis. The original report noted that only 6 out of 128 kinases have an IC_50_ estimated to be less than 100 nM, 51/128 kinases have an IC_50_ estimated to be between 100 nM and 1 µM, and 71/128 kinases have IC_50_ values estimated at greater than 1 µM. Careful analysis of the screening data suggests that Momelotinib will have an IC_50_ for Nek9 between 100 nM and 1 µM. Nek2 is the only other Nek with data reported in this publication and the Nek2 IC_50_ is estimated at > 1 µM. More recently, the selectivity of Momelotinib was determined using an affinity profiling method in a mixed cell lysate.^21^ These results indicated that Momelotinib inhibited Nek1, Nek3, and Nek9 with K_d_ values less than 100 nM. Finally, Momelotinib has been screened using the KINOMEscan technology of DiscoverX with the data set available via **L**ibrary of **I**ntegrated **N**etwork-based **C**ellular **S**ignatures (HMS LINCS data set ID 20082 found at http://lincs.hms.harvard.edu/kinomescan/). This screen at 10 µM implicates Nek3, Nek5, Nek6, Nek7, and Nek9 as potential targets, with little to no binding to Nek1, Nek2, Nek4, and Nek11. Momelotinib was not made to target the Nek family, so details of Nek structure activity relationships are unknown. Although a singleton result (one compound, no analogues), it is clear that this compound is a *bona fide* Nek inhibitor because three different assay formats all identified activity on one or more Nek kinases. Design of compound sets around this 2-anilino-4-aryl pyrimidine will likely lead to potent Nek inhibitors and improved understanding of Nek family SAR.

**Figure 3:**
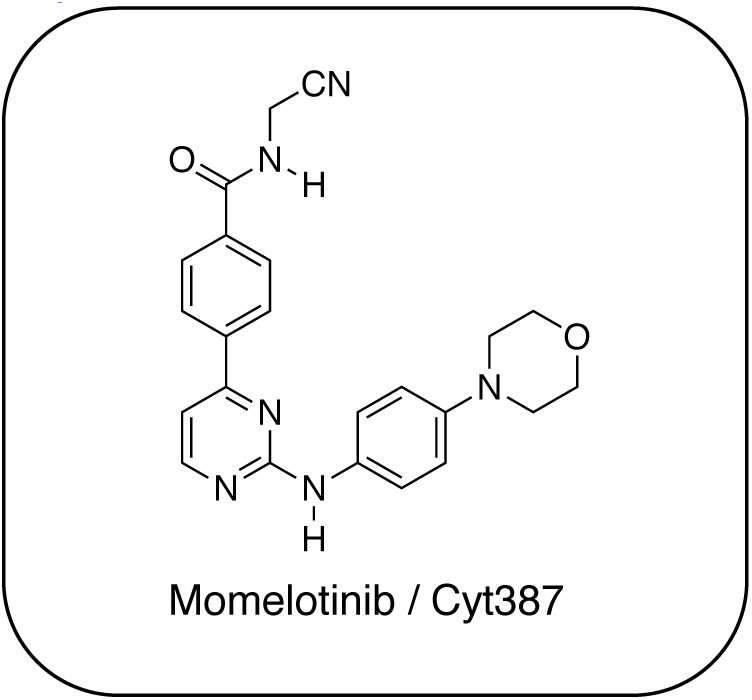
Momelotinib structure

Progress on understanding the intricacies of Nek biology will benefit from potent and selective Nek inhibitors. In addition, even the limited amount of work focused on the Nek family suggests that one or more of these understudied Nek kinases may emerge as a viable drug target. To date, there are few reports of medicinal chemistry efforts directed towards development of selective Nek kinase inhibitors. The work that has been done has focused on Nek2, and several Nek2 inhibitors with varying degrees of kinase profiling and selectivity have been published. Unfortunately, even these Nek2 inhibitors have not been tested systematically across the remainder of the Nek family. To design useful inhibitor tools for each specific Nek, we need to further understand the similarities and differences across the family to guide inhibitor synthesis. Here we address each Nek in turn and highlight potential chemical starting points that have emerged from our literature analysis. The vast majority of these compounds, which represent useful starting points for focused lead discovery efforts, arise from kinome cross screening, highlighting the value of collecting and sharing such data sets.

### Nek1

Nek1 has a functional role in the formation and regulation of cilia and its dysregulation is linked to several ciliopathies.^22–23^ Mutations in mouse Nek1 (*m*Nek1) resulted in progressive polycystic kidney disease where mice displayed facial dysmorphism, dwarfism, male sterility and anemia.^24^ *m*Nek1 was observed to localize to centrosomes during interphase and mitosis and is involved in cilia formation.^23^, ^25–26^ In humans, mutations in Nek1 have been linked to the lethal bone malformation disorder known as short-rib polydactyl syndrome type majewski.^27–29^ Nek1 is overexpressed in human gliomas suggesting it may be a potential therapeutic target for these brain tumors.^30^ Nek1 also regulates checkpoint control and DNA repair in response to DNA damage from UV or IR radiation, pointing to another potential application for Nek1 inhibitors in oncology.^31–33^

In addition to this basic research into Nek1 biology a crystal structure (PDB 4B9D) with a CDK inhibitor bound in the active site has been solved. Structures for apo- and ligand-bound Nek1 showed few differences with both adopting a “DFG-out/alpha-C out” inactive conformation. In contrast, both structures displayed the activation loop as ordered and in an extended conformation that is usually associated with an active kinase. On the other hand, the turn on the glycine-rich (P-) loop was disordered. Importantly, however, there have been no medicinal chemistry efforts undertaken to develop potent and selective Nek1 inhibitors. We examined the PKIS2 dataset to identify starting points for Nek1-directed chemistry. ^4^ The 4-aryl-7-azaindole scaffold (e.g. UNC5452) demonstrated Nek1 activity with only 18 off-target kinases (Table 5). Based on historical precedence for other kinases, SAR, and docking to the crystal structure, we propose that the azaindole makes two hydrogen bonds with the hinge region, and the sulfonamide group projects towards the catalytic lysine. (Figure 4a).

**Figure 4.**
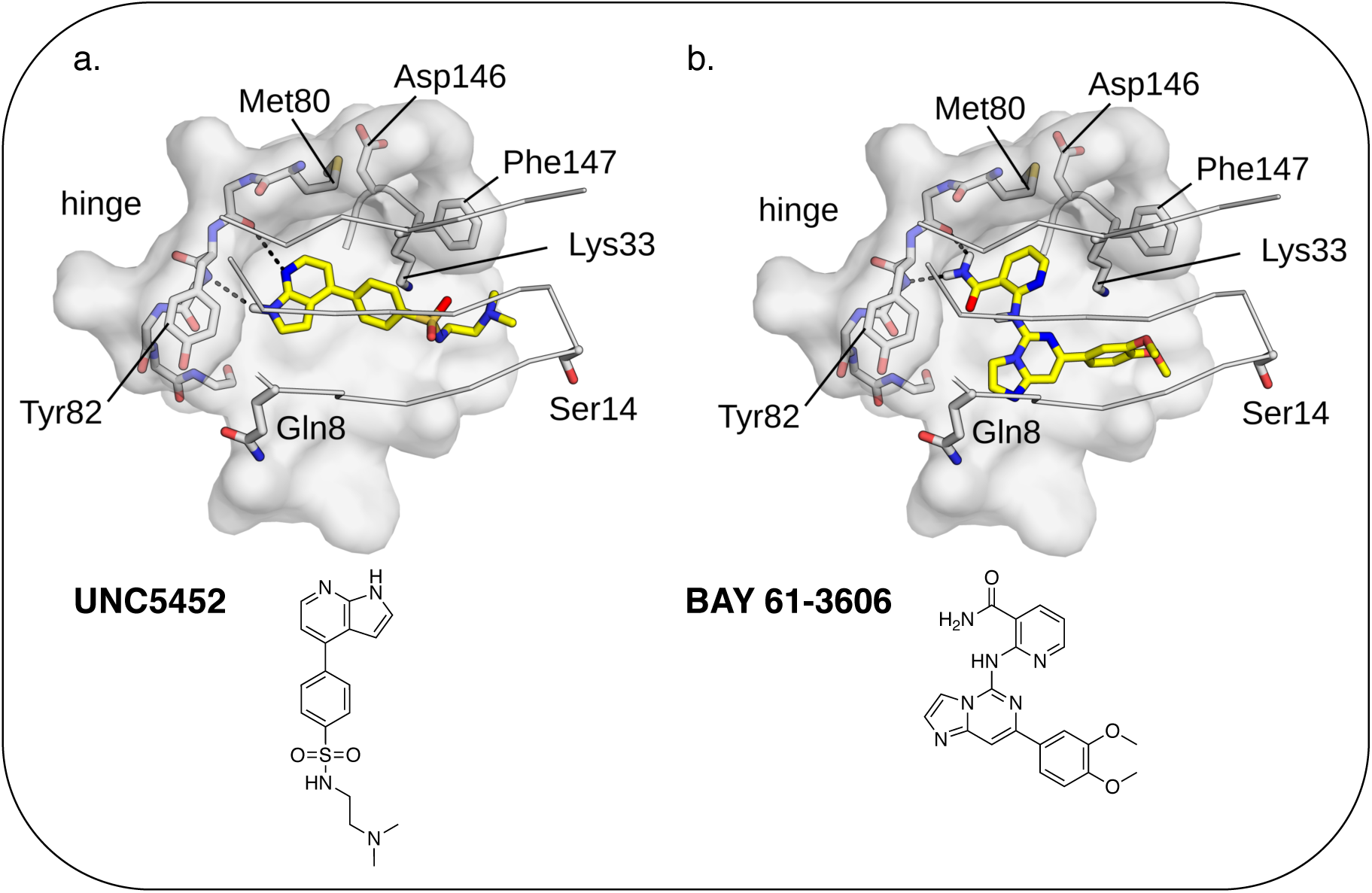
**a**. UNC5452 and **b**. BAY 61-3606 modeled into Nek1 crystal structure

**Table 5.**
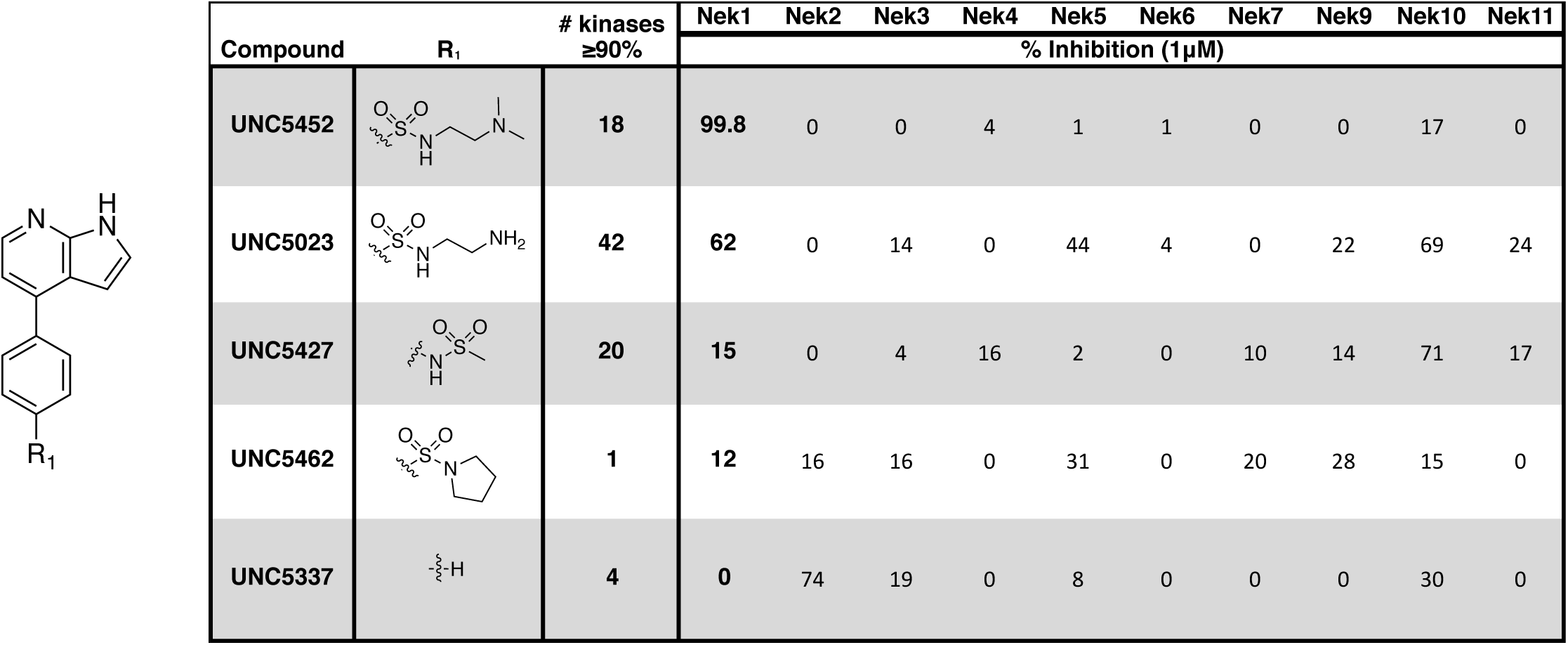
Percent inhibition values for the 4-aryl-7-azaindole PKIS2 series

| Compound | R<sub>1</sub> | # kinases<br>≥90% | Nek1 | Nek2 | Nek3 | Nek4 | Nek5 | Nek6 | Nek7 | Nek9 | Nek10 | Nek11 |
| --- | --- | --- | --- | --- | --- | --- | --- | --- | --- | --- | --- | --- |
| % Inhibition (1 μM) |  |  |  |  |  |  |  |  |  |  |  |  |
| UNC5452 |  | 18 | 99.8 | 0 | 0 | 4 | 1 | 1 | 0 | 0 | 17 | 0 |
| UNC5023 |  | 42 | 62 | 0 | 14 | 0 | 44 | 4 | 0 | 22 | 69 | 24 |
| UNC5427 |  | 20 | 15 | 0 | 4 | 16 | 2 | 0 | 10 | 14 | 71 | 17 |
| UNC5462 |  | 1 | 12 | 16 | 16 | 0 | 31 | 0 | 20 | 28 | 15 | 0 |
| UNC5337 |  | 4 | 0 | 74 | 19 | 0 | 8 | 0 | 0 | 0 | 30 | 0 |

The 4-aryl-7-azaindole, UNC5452, was originally published as an IKBKB inhibitor. There are five closely related compounds from this structural class in PKIS2 that demonstrate a range of potencies on Nek1 (Table 5). Removal of the terminal methyl groups of UNC5452 to give the primary amine UNC5023 resulted in a significant loss of binding. The three other analogues showed little to no binding.

The unusual combination of DFG-out/C-helix-out and an extended, activation loop, presents an opportunity for synthetic exploration to exploit these active site features in the design of selective, small-molecule kinase inhibitors. Based on our modeling, ligands that explore contacts with the P-loop could access two additional hydrogen bond donor/acceptors (Gln8 and Ser14). In both apo- and ligand-bound structures of Nek1, alpha-C is in the “out” conformation and is likely to be inaccessible to ATP-competitive small molecule inhibitors. Like most Nek-family members, Nek1 has an unusual Phe residue that can be targeted via the placement of ring systems between this residue and the gatekeeper (GK) methionine. Similar approaches have been successful in the design of Nek2 inhibitors. Additional compounds will need to be synthesized and evaluated for Nek1 potency and docked into this model to understand the SAR and the reasons for Nek1 selectivity.

The 4-thiophene-7-azaindole series is a related series identified in the PKIS2 dataset that showed a range of Nek1 activities (Table 6). This series is predicted to bind in the Nek1 model in the same orientation as the 4-aryl-7-azaindole but further work is required to fully understand the SAR and Nek selectivity profile.

**Table 6.** Percent inhibition values for 4-thiophene-7-azaindole series

| Compound | R1 | R2 | Nek1 % Inh at 1μM |
| --- | --- | --- | --- |
| UNC5078 |  |  | 91.9 |
| UNC5436 |  |  | 79 |
| UNC5265 |  |  | 74 |
| UNC5196 |  |  | 72 |
| UNC5503 |  |  | 71 |
| UNC5176 |  |  | 54 |
| UNC5041 |  |  | 48 |
| UNC5276 |  |  | 37 |
| UNC5108 |  |  | 15 |
| UNC5315 |  |  | 13 |
| UNC5471 |  |  | 0 |
| UNC5466 |  |  | 0 |
| UNC5412 |  |  | 0 |
| UNC5027 |  |  | 0 |

BAY 61-3606, a compound from the literature originally published as a SYK inhibitor,^34^ also showed potent Nek1 and Nek4 activity (159nM and 25nM respectively). ^35–36^ This compound was also docked into the Nek1 crystal structure with the amide moiety as the hinge binder (Figure 4b). This compound has two possible hinge binding groups based on literature precendent, the amide and the nitrogen of the 5-mambered ring of the imidazopyrimidine. Nek1 modeling suggested that the binding mode depicted in Figure 3b with the amide forming the hinge interaction is more likely because of possible steric clash with the catalytic lysine in the alternate binding mode. This particular binding mode is also supported by the lack of Nek1 activity observed with BAY 06-3606 analogues that removed the ortho amide.^37^

### Nek2

Nek2 is the most well studied member of the Nek kinase family. Nek2 is involved in key mitotic events and is localized at centrosomes in all stages of mitosis.^38^ Nek2 interacts with C-Nap1 (centrosomal Nek2-associate protein 1), PP1 (protein phosphatase 1), ninein-like protein (Nlp), and rootletin to regulate centrosome separation and microtubule organization.^39–42^ Nek2 also interacts with Mad1 (mitotic arrest deficient like-1), Mad2 (mitotic arrest deficient like-2), and Hec1 (highly expressed cancer 1) proteins to regulate spindle assembly checkpoint (SAC).^43–44^ Increased levels of Nek2 have been detected in several cancer types including breast carcinomas, hepatocellular carcinomas, and colon cancer, prompting the exploration of Nek2 inhibition for anticancer therapy.^45–48^

In concert with having a well-characterized biological role, Nek 2 has received the most attention in terms of small molecule medicinal chemistry efforts. There are a number of crystal structures of small molecules bound to Nek2 that can be used to facilitate further design. A list of all reported Nek crystal structures can be found in Supplemental Table 2. In 2009, GSK reported details of a thiophene benzimidazole series of PLK inhibitors.^49^ Cross-screening data from a small kinase panel for two advanced PLK compounds indicated that members of the series could exhibit a relatively narrow kinase inhibition profile. Interestingly, one of the off targets was Nek2. GSK compound 24 (Figure 5), for example, had an IC_50_ of 25 nM on Nek2. Out of 57 kinases screened only Nek2 and PLK had IC_50_ values below 100 nM, 7 kinases had IC_50_ values between 100 nM and 1 µM, and the remaining 48 kinases had IC_50_ values greater than 1 µM. A more advanced PLK1 compound in the same series, GSK461364A (Figure 5), was screened more broadly, and only inhibited 1.6% of the kinases across a large kinase panel with a potency less than 300 nM.^16^ In this assay, the K_d_ for Nek2 is 260 nM.

**Figure 5.**
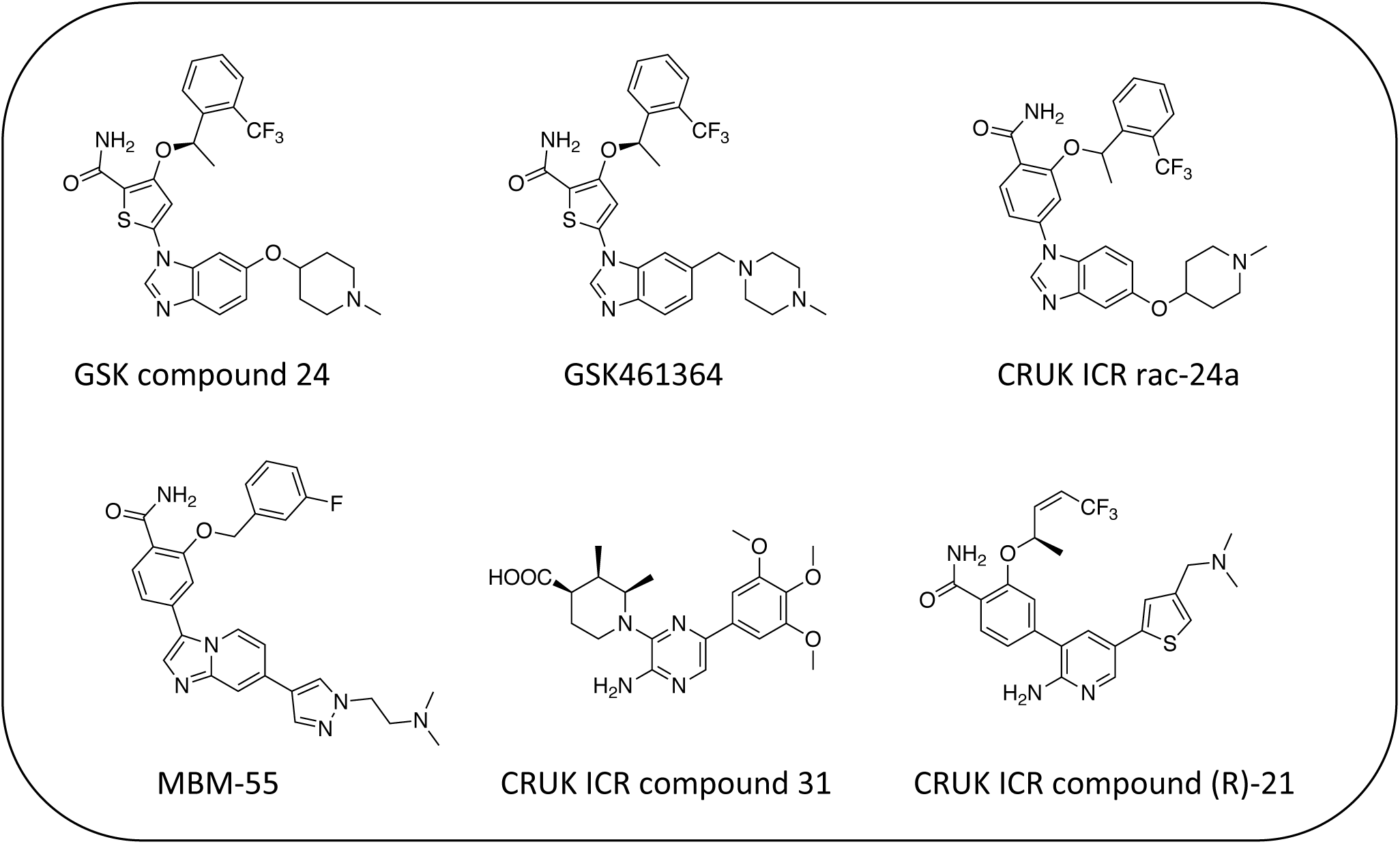
Examples of Nek2 inhibitors with benzimidazole, imidazopyridine, aminopyrazine, and aminopyridine hinge binding groups

A team of scientists at the Institute of Cancer Research utilized this same PLK series as a starting point for structure based design, with a primary aim being selectivity for Nek2 over PLK1.^50^ CRUK ICR Rac-24a (Figure 5) emerged from this work and has a Nek2 IC_50_ value of 570 nM and a PLK1 IC_50_ value of 88 µM. This compound demonstrates the ability to separate Nek2 activity from PLK1 activity. Two key changes that enhanced Nek2 selectivity were utilization of a phenyl ring as an isostere for the thiophene, and moving the phenoxy piperidine from the 6 position of the benzimidazole to the 5-position. The authors postulate that the different gatekeeper residues (Met for Nek2, Leu for PLK1) may account for selectivity improvements from the thiophene to phenyl switch. These aromatic rings pack closely with the gatekeeper residue. The phenoxy piperidine was moved to the 5-position because it was predicted this substituent would clash with Arg136 of PLK1, but Gly92 in this same position in Nek2 would be tolerant of such a substituent. Nek1, Nek2, Nek3, Nek4, Nek5, and Nek8 all have a methionine gatekeeper residue and a glycine residue that corresponds to the position of Gly92 in Nek2. It will be important to determine if this series shows activity on these other Nek family members, or if there is Nek2 selectivity based on other active site residue differences. Rac-24 was screened against a panel of 24 kinases (but no other members of the Nek family) and showed good selectivity, with only 4/24 kinases inhibited >50% when screened at 2 µM.

Recently new Nek2 inhibitors have been reported that replace the benzimidazole hinge binding moiety with the isosteric imidazo[1,2-a]pyridine ring system.^51^ This team identified the most potent Nek2 inhibitors currently in the literature. Compound MBM-55 (Figure 5), for example, had a Nek2 IC_50_ value of 1 nM. MBM-55 was screened against 12 other kinases. Five of these kinases had IC_50_ values under 100 nM (RPS6KA1 = 5.4 nM, DYRK1A = 6.5 nM, ABL1 = 20 nM, CHEK1 = 57nM, and GSK3B = 91nM). In order to interpret the biological activity of these compounds it will be important to determine their potency on the remainder of the Nek family and across the whole kinome.

CRUK ICR compound 31 (Figure 5) is representative of the aminopyrazine class of Nek2 kinase inhibitors, which was optimized by structure-based design effort from a screening hit.^52^ This compound has a Nek2 IC_50_ value of 230 nM, and a Nek1 IC_50_ value of 170 nM. The compound shows selectivity over PLK1 (IC_50_ = 19 µM) and CHEK1 (IC_50_ > 100 µM), but broad kinome profiling data (including other Nek family members) on this series is not available. The binding mode of this class was determined by X-ray crystallography. An important finding was that the compounds bound to an unusual inactive kinase conformation in which Tyr70 moved into the active site and interacted with the carboxylic acid of the inhibitor. The authors hypothesized that the rare combination of Phe148 and a large gatekeeper residue, introduces steric constraints in the ATP binding site, perhaps making it difficult to target with small molecules.

An aminopyridine series was designed for Nek2 by combining elements of the prior aminopyrazine series SAR and the benzimidazole series SAR.^53^ This work culminated in the identification of compound CRUK ICR (R)-21 (Figure 5), with Nek2 IC_50_ = 22 nM, and PLK1 IC_50_ = 5.8 µM. (R)-21 was screened against 24 additional kinases (but no additional Nek family members). The IC_50_ for GSK3B inhibition was 70 nM, and 110 nM for LCK inhibition. Four additional kinases had IC_50_ values between 300 nM and 1 µM, and the remaining 18 kinases had potencies > 1 µM.

Scientists at UCSF took advantage of a rare active site cysteine in Nek2 to design irreversible inhibitors that show cellular activity.^54^ Compound JH295 (Figure 6), an oxindole with an electrophilic propynamide, showed the best balance between potency, selectivity, and reactivity. The ethyl group on the imidazole ring was carefully chosen to enhance selectivity, for example over CDK1, a key off target whose inhibition would confound interpretation of cellular results.

**Figure 6:**
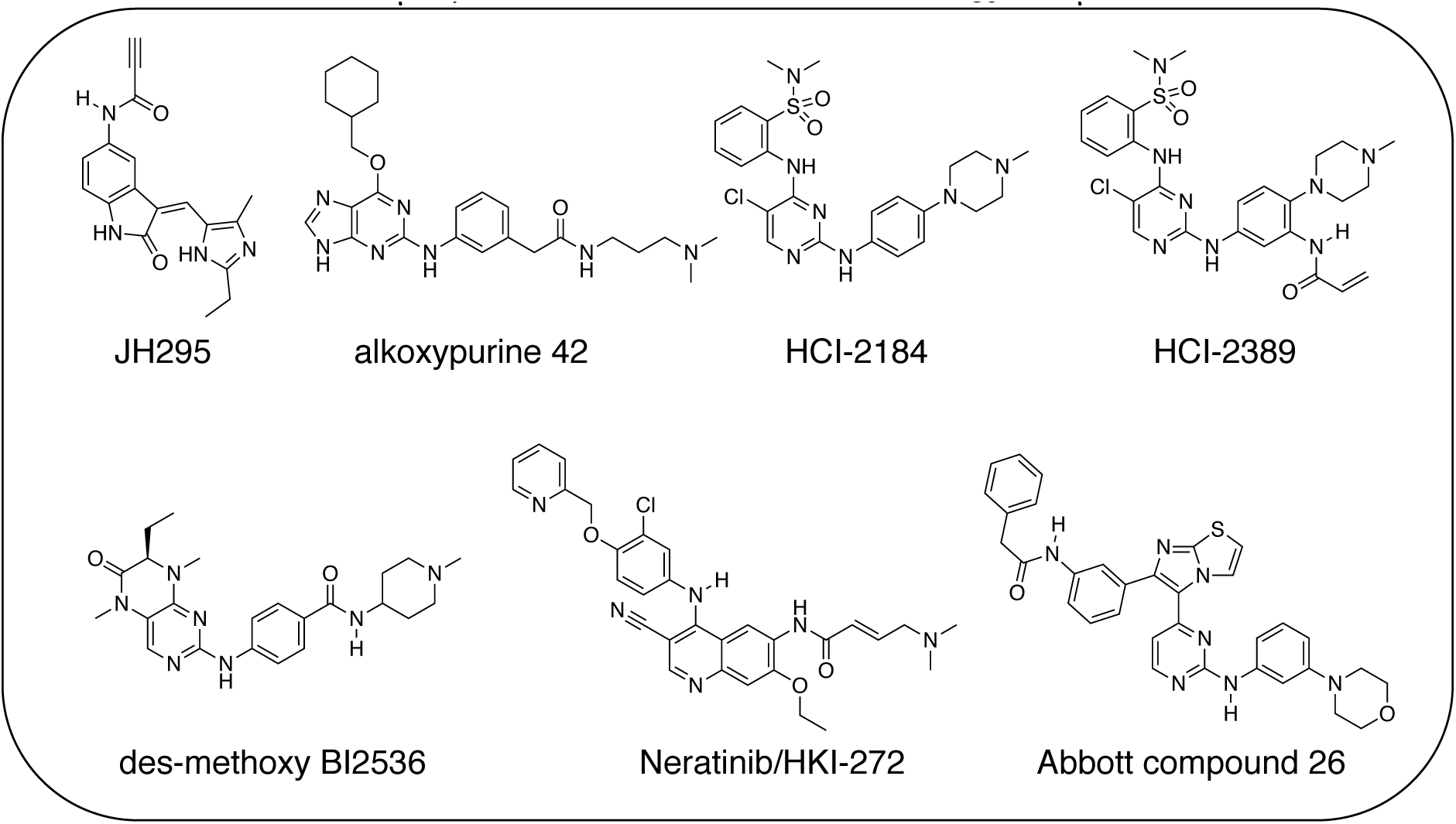
Nek2 inhibitors with a variety of hinge binding groups

Alkoxypurine 42 (Figure 6) inhibits Nek2 with an IC_50_ value of 280 nM.^55^ It showed promising selectivity over CDK2 (IC_50_ = 1 µM). X-ray crystal structures of members of the series in complex with Nek2 showed 3 hydrogen bonds to the hinge region of the kinase. The pendant aniline group points towards the solvent front. The cyclohexyl moiety projects towards the catalytic lysine and the ribose binding pocket. Both points of diversity could likely be explored further to build in potency and selectivity. No results were reported for other Nek family members. This scaffold, with robust chemistry and a well-defined binding mode, could be further explored to identify inhibitors of other Nek family members as well.

Scientists researching therapies for multiple myeloma discovered that high Nek2 expression correlated with drug resistance in this incurable disease. This result prompted them to look for Nek2 inhibitors that could perhaps play a role in the treatment of multiple myeloma.^56^ They identified pyrimidine HCI-2184 (Figure 6), a reversible inhibitor, and pyrimidine HCI-2389 (Figure 6), an irreversible analogue of HCI-2184 with an electrophilic acrylamide group. Both HCI-2184 and HCI-2389 were tested against 451 kinases. Data was only reported for 39 representative kinases. These results and broad kinome screening of closely related dianilinopyrimidines such as TAE-684 suggest that the compounds may be active on too many kinases to ascribe the biological consequences solely to Nek2 inhibition. Nevertheless, this scaffold may prove suitable for optimization for Nek2 or Nek1 activity. Nek1 is the only other Nek for which data is available.

PLK1 and Nek2 share some common features of their active sites, including the rare combination of a large gatekeeper residue (Met86 in Nek2 and Leu130 in PLK1) and a phenylalanine forming a lipophilic surface underneath the adenine ring of ATP (Phe148 in Nek2, Phe183 in PLK1)^57^, so it is not surprising that there is chemical matter that cross reacts with these two kinases. In addition to the aforementioned thiophene benzimidazole series, there is evidence of some cross reactivity in a pyrimidine pteridonone series of PLK1 inhibitors as well. BI2536 is a relatively selective PLK1 inhibitor. Removal of the methoxy group on the aniline that points towards solvent gives des-methoxy BI2536 (Figure 6), and this compound gives 72% inhibition of Nek2 activity at a concentration of 1 µM.^57^ The methoxy group of BI2536 fits into a small lipophilic pocket near the hinge, adjacent to Leu132. Nek2 has a tyrosine in this position that would sterically clash with the methoxy group. Elimination of the methoxy removes the steric mismatch adjacent to the hinge and allows for Nek2 inhibition.

A team studying Nek2 overexpression in *Drosophila melanogaster* in order to elucidate contributions of Nek2 to tumorigenesis identified the EGFR inhibitor HKI-272 (Figure 6),^58^ also known as Neratinib, as a Nek2 inhibitor with a potency of 274 nM.^59^ Neratinib has been screened in a broad panel of kinases, and showed activity on 9.3% of the kinases screened with a K_d_ of less than 300 nM.^16^ Broad screening results are also available through the HMS LINCS database (dataset ID 20053 and dataset ID 20195), where 34 kinases show a K_d_ of less than 250 nM. In this assay format, the Nek2 K_d_ is 270 nM, which compares favorably with the 274 nM value determined in the enzyme assay. Although Neratinib is not exquisitely selective, it represents a useful starting point for synthesis of potent and selective Nek2 inhibitors given its relatively narrow kinase profile.

A compound from a very different chemotype emerged from an SAR study on imidazothiazoles designed to simultaneously target the kinases IGF1R, EGFR, and ErbB2.^60^ These compounds differ from all other Nek2 inhibitors described here because they are much larger and have a lipophilic acetamide group that projects into the back pocket of kinases like ErbB2. Abbott compound 26 (Figure 6) was screened against a total of 59 kinases. It had an IC_50_ of 310 nM against Nek2, the only Nek screened. Overall, it inhibited 9/59 kinases with an IC_50_ under 100 nM, 10/59 with an IC_50_ between 100 nM and 1 µM, and 40/59 kinases with an IC_50_ > 1 µM.

Tables 7, 8, and 9 present data for a family of molecules that exhibit cross reactivity with Nek2. These results can guide medicinal chemistry efforts in the pursuit of Nek2 selective compounds. The Nek2 SAR available in the literature is organized in Tables 7, 8, and 9. The compounds in Table 7 originate from a GSK project to develop selective EIF2AK3 (PERK) inhibitors.^61^ These compounds are part of PKIS2 and were screened against a large kinase panel.^4^ Table 7 summarizes the Nek family results.

**Table 7.** Nek2 SAR for a series of published PERK inhibitors

| Compound | R1 | # of kinases<br>with %Inh >90 | NEK1 | NEK2 | NEK3 | NEK4 | NEK5 | NEK6 | NEK7 | NEK9 | NEK10 | NEK11 |
| --- | --- | --- | --- | --- | --- | --- | --- | --- | --- | --- | --- | --- |
|  |  |  | %Inh (1µM) |  |  |  |  |  |  |  |  |  |
| UNC5254 |  | 15 | 89 | **97** | 0 | 29 | 34 | 0 | 0 | 0 | 0 | 58 |
| UNC5490 |  | 16 | 31 | **96.9** | 7 | 11 | 45 | 19 | 0 | 20 | 0 | 59 |
| UNC5071 |  | 11 | 7 | **84** | 0 | 32 | 8 | 18 | 0 | 8 | 10 | 38 |
| UNC5172 |  | 86 | 0 | **79** | 0 | 5 | 7 | 3 | 17 | 33 | 0 | 0 |
| UNC4987 |  | 15 | 21 | **74** | 4 | 44 | 24 | 18 | 11 | 25 | 7 | 49 |
| UNC5297 |  | 14 | 31 | **73** | 11 | 22 | 28 | 0 | 0 | 0 | 22 | 8 |
| UNC5039 |  | 17 | 26 | **66** | 1 | 22 | 32 | 11 | 0 | 18 | 18 | 0 |
| UNC5478 |  | 13 | 9 | **18** | 0 | 13 | 0 | 6 | 0 | 25 | 6 | 8 |
| UNC5165 |  | 5 | 0 | **15** | 0 | 7 | 0 | 0 | 11 | 8 | 18 | 14 |

All of the compounds in Table 7 are potent PERK inhibitors (data not shown). UNC9054 was the weakest PERK inhibitor, with a still potent IC_50_ = 17 nM. The remaining compounds had PERK IC_50_ values below 1 nM. Inhibition of Nek2 was greatly reduced when the R_1_ group had a CF3 group in the meta position (UNC5478 and UNC5165). UNC5490 showed 97% inhibition of Nek2 binding at a screening concentration of 1 µM in a DiscoverX panel with selectivity over the other Nek kinases. Nek SAR for a related set of compounds with some core and side chain modifications is shown in Table 8. These come from the same PERK project at GSK and the compounds are included in PKIS2. All the compounds in Table 8 have PERK IC_50_ values below 15 nM, but are also selective for Nek2 within the Nek family. Nek2 tolerates the addition of the pyrazole ring in UNC5500, a region of the active site that is unexplored in the compounds of Table 7. This compound is an inhibitor of 36 kinases (> 90% Inh at 1 µM), however, so it would need additional optimization to remove the off-target activity in a Nek2 optimization effort.

**Table 8.** Nek2 SAR for a series of PERK inhibitors

| Compound | X | Z | R1 | R2 | # of kinases with %I>90 | NEK1 | NEK2 | NEK3 | NEK4 | NEK5 | NEK6 | NEK7 | NEK9 | NEK10 | NEK11 |
| --- | --- | --- | --- | --- | --- | --- | --- | --- | --- | --- | --- | --- | --- | --- | --- |
| %Inhibition (1μM) |  |  |  |  |  |  |  |  |  |  |  |  |  |  |  |
| UNC5500 | S | C |  |  | 36 | 7 | 85 | 0 | 11 | 21 | 3 | 0 | 0 | 22 | 87 |
| UNC5000 | O | N | none |  | 2 | 0 | 70 | 0 | 0 | 22 | 0 | 0 | 4 | 1 | 2 |
| UNC5369 | S | C |  |  | 22 | 0 | 70 | 1 | 17 | 22 | 8 | 19 | 28 | 14 | 44 |
| UNC5012 | S | C |  |  | 37 | 0 | 28 | 1 | 0 | 5 | 0 | 0 | 0 | 22 | 47 |
| UNC5494 | S | C | H |  | 1 | 0 | 7 | 0 | 9 | 6 | 0 | 0 | 4 | 12 | 9 |
| UNC5119 | S | C | H |  | 2 | 0 | 4 | 10 | 0 | 14 | 0 | 0 | 18 | 9 | 0 |

Table 9 shows results for three additional compounds (Abbott 405, Abbott 502, and Abbott 511) described in the supplemental material of a paper that described broad kinase screening for a large set of molecules.^36^ A broad range of chemotypes was tested in this exercise, and importantly, there is often data for multiple closely related molecules from the same chemical clusters. The identification of these three Nek2 inhibitors, which are quite similar to the compounds in Tables 7 and 8, by a different research group using a different assay format gives confidence that this chemotype represents a robust starting point for synthesis of Nek2 inhibitors. Of the three compounds, Abbott 502 seems to be the superior Nek hit because it only has activity towards 12 percent of the kinases screened with an IC_50_ < 300 nM. Abbott 405 and Abbott 511 are more promiscuous, with S(300 nM) scores of 0.44 and 0.39, respectively.

**Table 9.** Nek2 SAR from Abbott broad kinase screening

| Compound | R1 | R2 | S(300 nM) | NEK2 | NEK4 |
| --- | --- | --- | --- | --- | --- |
|  |  |  |  | IC50 (nM) |  |
| Abbott 405 |  |  | 0.44 | 13 | 200 |
| Abbott 502 |  |  | 0.12 | 40 | 1000 |
| Abbott 511 |  |  | 0.39 | 50 | >3900 |

### Nek3

Nek3 is a cytoplasmic Nek kinase first cloned and characterized in 1999.^62^ Not many cellular functions of Nek3 are currently known, except its association with the human breast cancer hormone prolactin (PRL). By regulating the phosphorylation of Vav2 (a nucleotide exchange factor that is a part of PRL signaling) Nek2 plays a role in PRL mediated effects on human breast cancer survival, motility and progression. This role of Nek3 was further substantiated when T47D cells transfected with kinase dead Nek3 displayed enhanced apoptosis than normal cells.^63^ In a separate study, overexpression of Nek3 in Chinese Hamster Ovary (CHO) cells led to enhanced PRL mediated cytoskeletal rearrangement whereas down regulation of Nek3 showed the opposite effect.^64^ Threonine 165 is a key regulatory site for the activation of Nek3. Expression of a mutant Nek3 (Nek3-T165V) that cannot be phosphorylated at this spot on the activation loop into MCF-7 cells resulted in cells with increased focal adhesion size and reduced propensity to migrate. ^65^ This clearly shows the role of Nek3 in enhancing breast cancer migration and invasion, and that inhibition of Nek3 is an attractive approach for breast cancer therapy. Nek3 is highly expressed in post mitotic neurons of mouse brain, especially during embryogenesis and post-natal life indicating its role in early brain development. Nek3 mutations resulted in microtubule deacetylation, thus suggesting a potential role of Nek3 in disorders like Huntington’s disease where microtubule function is impaired.^66^ Nek3, along with 16 other human protein kinases, is also an essential host kinase required for influenza virus replication.^67^

The ability to inhibit Nek3 potently and selectively is potentially important to understanding the preliminary research linking Nek3 inhibition to diseases such as breast cancer and Huntington’s disease. Based on our analysis of the literature there were no compounds specifically designed for Nek3. One series of compounds that we found in the literature was the amino pyrimidine compounds JNK-IN-7^68^ and JNK-9L^69^ (Figure 7). Both of these compounds were originally designed as inhibitors for JNK1/2/3 (Gene names: MAPK8, MAPK9, MAPK10 respectively). JNK-IN-7 has IC_50_ values of 41, 25 and 13 nM for JNK1, JNK2, and JNK3 respectively. The binding pose of compound JNK-IN-7 as revealed from a cocrystal structure with JNK3 shows that the amino pyrimidine core is involved in a hydrogen bond interaction with the hinge and the acrylamide moiety is involved in a covalent bond formation with a cysteine residue. A complete kinome scan of compound JNK-IN-7 revealed that in addition to JNK1/2/3 activity, it showed more than 90% inhibition of Nek3, Nek5 and Nek7 kinase activity (Table 10). It should be noted that close structural analogs JNK-IN-1, JNK-IN-8, JNK-IN-11 and JNK-IN-12 (Figure 7), also possessing an electrophilic acrylamide moiety, did not show any appreciable inhibition of Nek3. The lack of affinity for Nek3 of JNK-IN-1 and JNK-IN-8 suggests that the methyl group in these compounds might offer steric blockade or conformational modification that prevents suitable interaction with the Nek3 active site. A possible explanation is the presence of a tyrosine residue at the hinge of Nek3, which may make it too crowded for the ortho-methyl group. This design strategy to improve selectivity has been successfully applied for other kinases. Similarly in the case of JNK-IN-11 and JNK-IN-12 the bulky substituents on the pyrimidine ring might lead to an unsuitable orientation of these compounds, resulting in the loss of affinity. JNK-9L, which lacks the electrophilic acrylamide moiety, also displayed Nek3 inhibition, thus suggesting that Nek3 inhibition by these compounds was not necessarily due to covalent interaction. Additional analogs would need to be synthesized and screened to evaluate the SAR around this series and optimize for Nek3 activity and away from the JNK family.

**Table 10.**
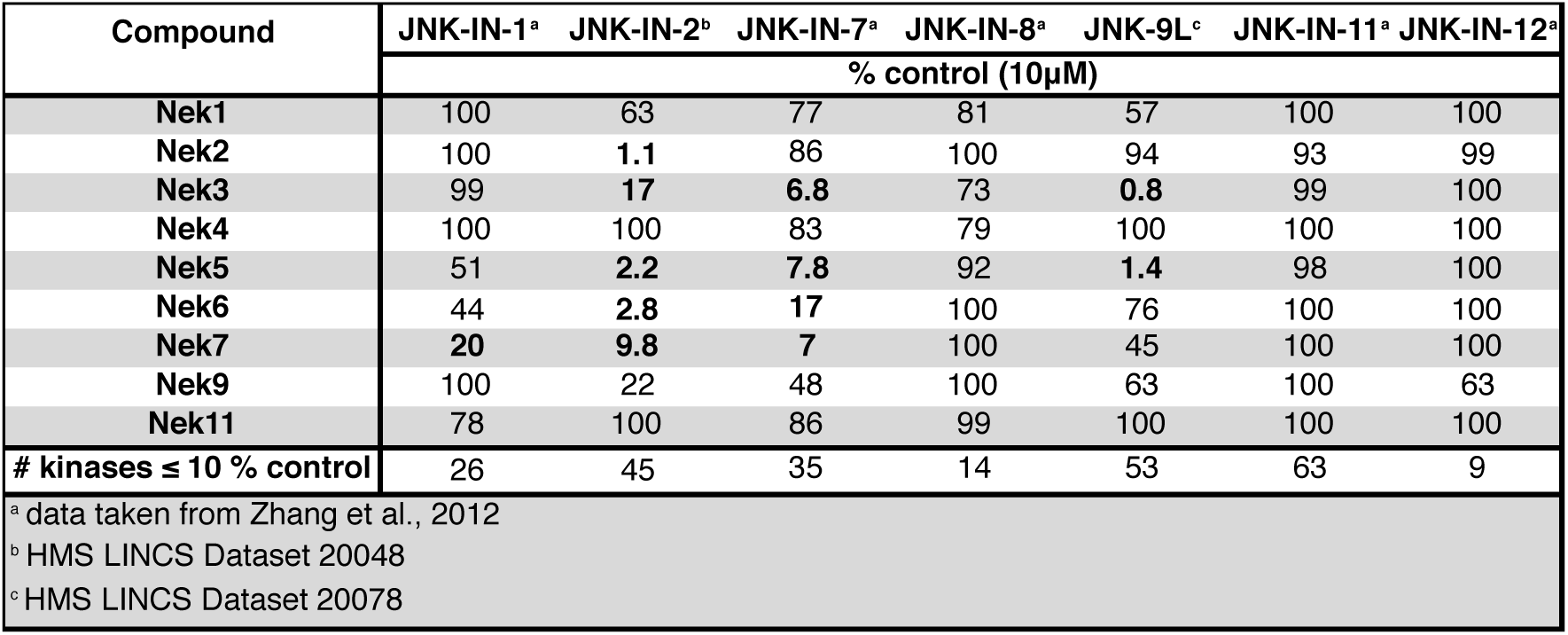
Selectivity profile of JNK inhibitors across the Nek family

**Figure 7.**
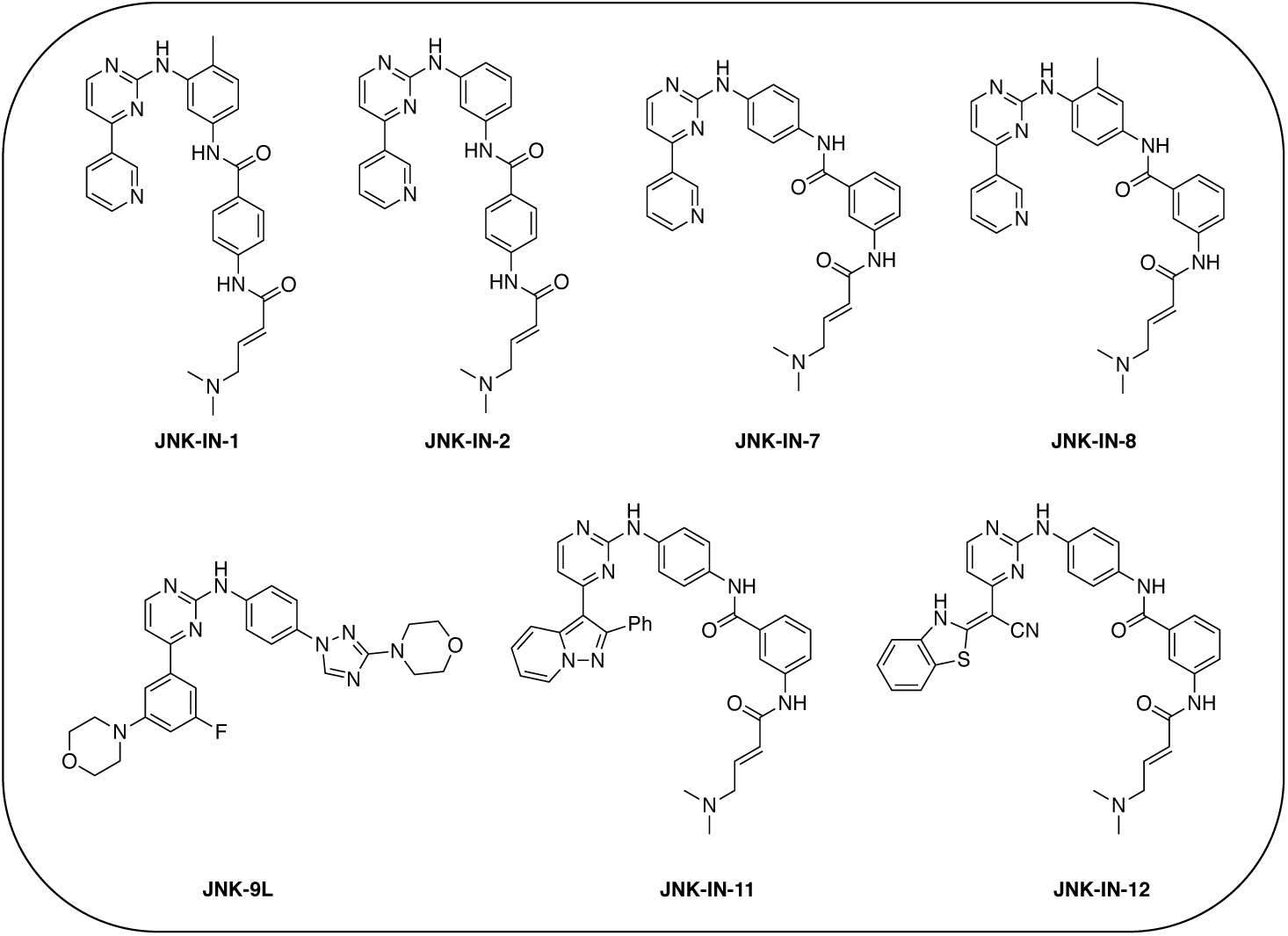
Structures of JNK inhibitors, some of which have Nek cross reactivity

### Nek4

In 2010 Nek4 kinase was identified as a mediator of cellular responses to microtubule disrupting drugs. The interplay between Nek4 activity and cells’ responses to various microtubule-disrupting agents is not straightforward. For example, when human lung adenocarcinoma cells were treated with the microtubule stabilizer Taxol, Nek4 deficiency led to resistance. In contrast, Nek4 deficiency in cells treated with the microtubule-disrupting agent Vincristine resulted in increased sensitivity to Vincristine. Thus Nek4 deficiency is involved in microtubule homeostasis and either antagonizes or enhances the effect of drugs targeting microtubules.^70^ Suppression of Nek4 in human fibroblast cells resulted in impaired cell cycle arrest in response to double stranded DNA damage initiated by Etoposide and Mitomycin C.^71^ Nek4 is highly upregulated in lung and colon cancer and is associated with the apoptosis triggering *tumor necrosis factor (TNF)-related apoptosis-inducing ligand (TRAIL)* in lung cancers.^72^ A possible role of Nek2/Nek4 in zygote to ookinete transition in plasmodium parasites was recently shown.^73^

Due to the limited details of Nek4 biology, a potent and selective small molecule Nek4 inhibitor would be beneficial in determining the consequences of Nek4 inhibition and its role in disease. Towards this goal, we have identified several Nek4 inhibitor chemical starting points. These compounds were not synthesized as Nek4 inhibitors, but Nek4 activity was observed as an off-target, and this cross reactivity can be exploited. For example, BAY 61-3606 (Figure 8) was originally developed as an orally available, selective SYK inhibitor^34^ but was later reported to have both Nek1^37^ and Nek4^36^ activity. BAY 61-3606 has two potential hinge-binding moieties - the amide or the nitrogen at the 1-position of the imidazopyrimidine. Additional analogues of this compound have been profiled against Nek1 and Nek4, but lack activity (Figure 9).^37^ This result lends credence to the idea that the amide in the ortho position may be the hinge-binding group and is required for Nek1 and Nek4 activity. Further SAR or crystallographic confirmation will be required to confirm this theory.

**Figure 8.**
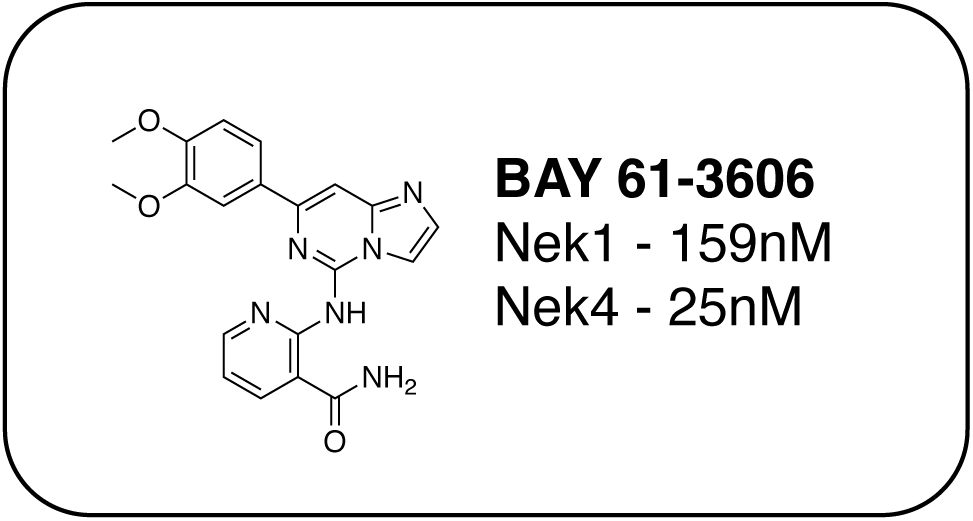
Structure of BAY 61-3606

**Figure 9.**
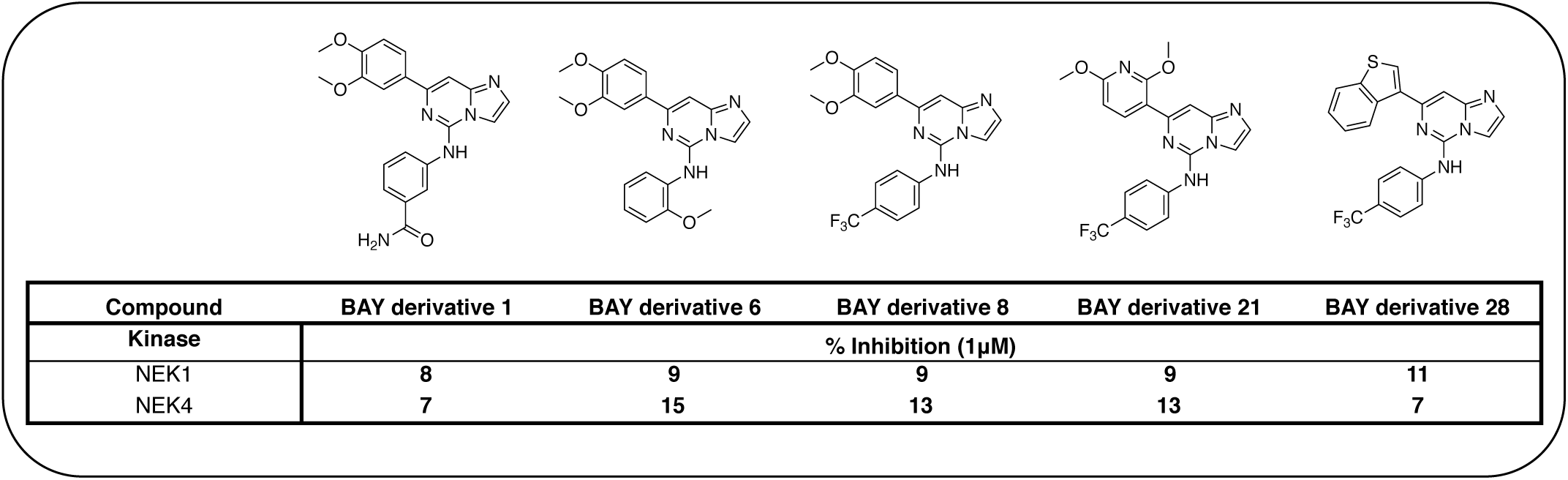
Structural analogues of BAY 61-3606 with and Nek1 and Nek4 activities

A thienopyridine series that we have identified from the literature is a possible starting point to obtain a potent and selective Nek4 inhibitor (Table 11). Abbott Laboratories originally described Abbott compound 17 as a MAP3K8 (COT) inhibitor.^74^ The cross-screening for this compound against many kinases revealed activity on Nek4.^36^ Two other compounds (Abbott compounds 3359 and 3357) from this series also inhibited Nek4, although none were as potent as Abbott compound 17 (Table 11). No member of this series inhibited Nek2 (IC_50_ >10,000nM). This series was further elaborated and repurposed by Genentech, resulting in development of potent CDK8 inhibitors.

**Table 11.** SAR for thienopyrimidine series

| Compound | R<sub>1</sub> | R<sub>2</sub> | Nek4 IC<sub>50</sub> nM | Nek4 %Inh | COT IC<sub>50</sub> (nM) |
| --- | --- | --- | --- | --- | --- |
| Abbott Cmpd 17 |  |  | 50 | — | 1120 |
| Genentech Cmpd 1 |  |  | — | 47.5<sup>a</sup> | 410 |
| Genentech Cmpd 2 |  |  | — | 89.9<sup>a</sup> | 150 |
| Genentech Cmpd 32 |  |  | — | 8.7<sup>b</sup> | — |
| Abbott Cmpd 3359 |  |  | 316 | — | — |
| Abbott Cmpd 3357 |  |  | 1000 | — | — |

<sup>a</sup> 100nM screening concentration
<sup>b</sup> 1 μM screening concentration

Notably some Nek4 activity emerged from the cross screening of these COT and CDK8 inhibitors. For example, replacement of oxadiazoline with a tetrazole was tolerated when R_2_ was a substituted phenyl (Genentech Compound 2). When the halogen was maintained at R_2_ and the heterocycle was replaced by an amide (Genentech Compound 32) the analogue lost Nek4 activity.^75^ Genentech compounds 1 and 2 were not counter-screened against Nek2 but were screened in a panel of 51 kinases. They only inhibited 6 and 4 kinases, respectively, above 80% inhibition at a screening concentration of 100 nM.

A third potential Nek4 series is shown in Figure 10. Abbott Compound 1141 is a simple biphenyl structure that employs pyridine as a hinge binding group. It has sub-100 nM potency on Nek4, and inhibits 15/131 kinases with IC_50_ values < 300 nM. This chemotype lends itself to an efficient modular synthetic approach to elucidate Nek4 SAR using robust chemistry and readily available building blocks. Synthesis of additional analogues would seek to improve the Nek4 potency, while reducing activity on kinases that are potently inhibited by Abbott 1141, such as ROCK1, PRKG1, and PRKG2.

**Figure 10.**
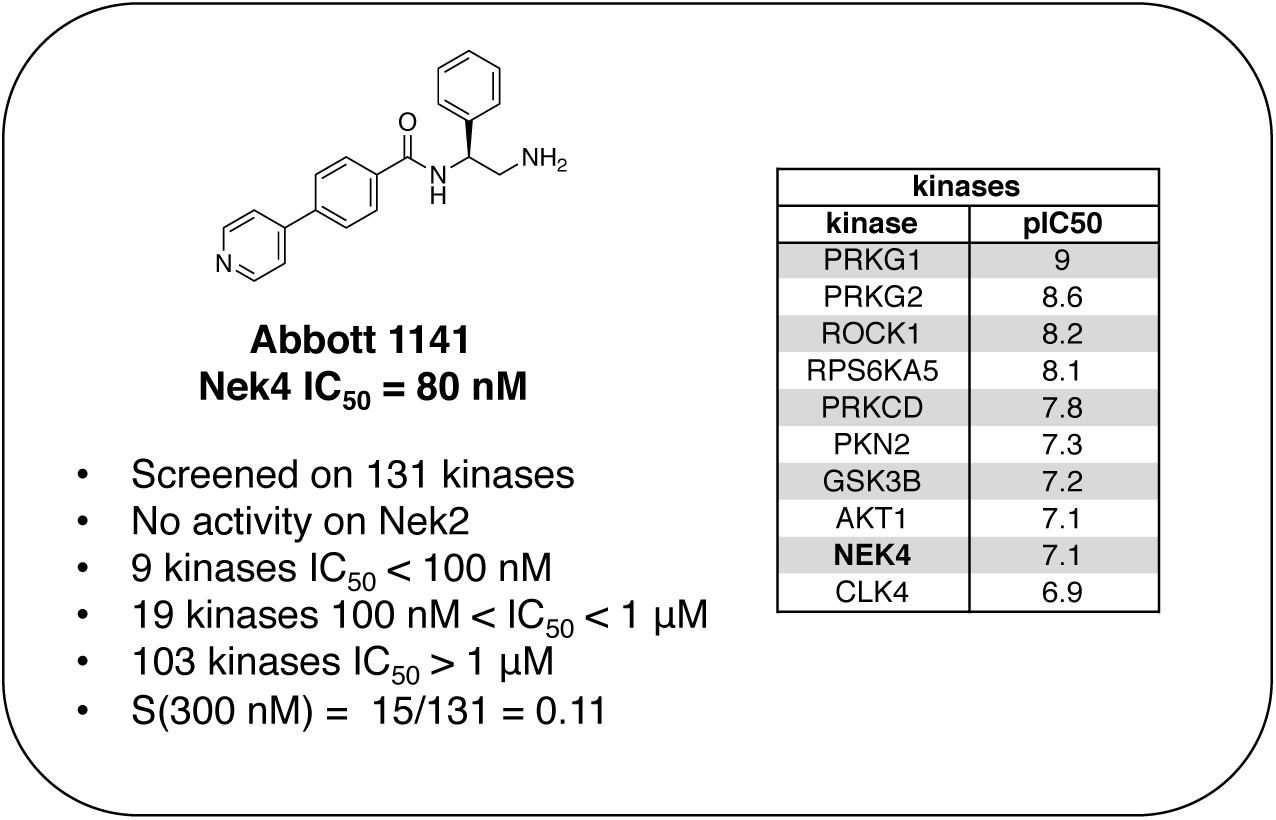
Abbott 1141 structure and activity profile

These structurally different scaffolds can provide starting points for generation of potent Nek4 inhibitors with a narrow kinase inhibition profile. Currently Bay 61-3606 has good Nek4 potency, but inhibits too many additional kinases (14/385 >90% Inhibition at 1µM) to be used to elucidate Nek4 specific biology. Abbott Compound 17 and 1141 also have very good Nek4 potency, but inhibited 12/137 and 15/131 kinases respectively greater than 300 nM. Further screening on both of these compounds would be required to determine their selectivity against the other Nek family members and broader kinome wide selectivity.

### Nek5

An extremely dark member of the Nek kinase family, Nek5 was recently shown to be involved in the regulation of centrosome integrity. Similarly to Nek2, depletion of Nek5 resulted in delayed centrosome separation leading to reduced microtubule nucleation and errors in chromosome segregation.^76–77^ In a separate study, expression of Nek5 in Hek293 cells resulted in enhanced cell viability and reduced cell death, whereas depletion of Nek5 reduced cancer cell drug resistance and induced apoptosis in vitro.^78^

Nek5 is the kinase in the Nek family with the least information known about its function and subcellular localization.^78^ Excitingly, however, the PKIS2 dataset identified multiple possible chemical starting points for a Nek5 inhibitor. Remarkably, Nek5 had the highest hit rate (0.065) across all of the Nek kinases that were screened (Table 2). The 2-anilino-4-pyrrolidinopyrimidines have 39 exemplars that showed selectivity within the Nek family, and did not inhibit any other Nek family members greater than 68%. The series had K_d_ measurements on Nek5 ranging from 370 nM to 2300 nM. Based upon these results and the number of compounds that were screened some structural requirements for activity can be identified. A change of the trimethoxy aniline to a para fluoro aniline at the R_1_ position was not tolerated and results in a total loss of potency (Table 12). Modification of the amide to the urea also caused a decrease in potency. Interestingly the pyrolidine amide could be swapped for a piperidine sulfonamide (UNC5124) without a loss in potency. This series is both relatively Nek family specific and also kinome selective with the most potent analog only inhibiting 3 other kinases above 90% at 1 µM. Further analogues aimed at exploring the aniline in the 2 position, ring size, stereochemistry, and nitrogen capping groups should be straightforward to synthesize and would elucidate the SAR for Nek5.

**Table 12.** Percent Inhibition and K_d_ values of 2-anilino-4-pyrrolidinopyrimidines

| Compound | n | R<sub>1</sub> | R<sub>2</sub> | % Inh (1μM) | K<sub>d</sub> (nM) | # kinases ≥90% |
| --- | --- | --- | --- | --- | --- | --- |
| UNC5052 | 1 |  |  | 97.3 | 370 | 4 |
| UNC5135 | 1 |  |  | 99.9 | 460 | 6 |
| UNC5391 | 1 |  |  | 93.1 | 730 | 7 |
| UNC5112 | 1 |  |  | 35 | – | 0 |
| UNC5124 | 2 |  |  | 100 | – | 13 |

A series of JNK inhibitors from the literature have also been shown to inhibit Nek5. Two of these compounds, JNK-IN-2 and JNK-IN-7 (Figure 7), inhibit Nek5 at 2.2 and 7.8 percent of control, respectively (Table 10). Both of these compounds contain an acrylamide that is capable of forming a covalent bond with a cysteine. In JNK kinases the cysteine that is utilized for covalent binding is in the hinge binding region.^79^ Nek5 also has a cysteine available in the hinge region however it is transposed by four amino acids. If this cysteine were capable of being targeted by a reactive group to form a covalent bond it would be a viable design strategy to improve selectivity and potency. To validate whether Nek5 was capable of being targeted by an irreversible inhibitor further experimental validation would need to be undertaken. A third JNK inhibitor that also inhibits Nek5 is JNK-9L (Figure 7). This compound has a similar shape to JNK-IN-2 and JNK-IN-7 but does not contain a moiety capable of forming a covalent bond. This compound, although fairly Nek family selective, also inhibited Nek3, and importantly inhibited 53 kinases below 10% control and is likely too promiscuous to be a good chemical starting point.

### Nek6

Nek6 kinase is phosphorylated and activated during the M phase and is required for the progression of mitosis. Inhibiting Nek6 function led to cell arrest in M phase and induced apoptosis.^80^ Nek6 also phosphorylates Eg5, a kinesin protein that is critical for mitotic spindle formation.^81^ Nek6 is phosphorylated by the checkpoint kinases CHEK1 and CHEK2, highlighting its role in cell cycle checkpoint control in response to DNA damage.^82^ Nek6 is also upregulated in a wide range of malignancies, including breast cancer, colon cancer, lung cancer, gastric cancer, melanoma, non-Hodgkins lymphoma, and glioblastoma.^83–85^

Although there has been one report on virtual screening for Nek6^86^, and one report on a thermal shift assay for several Nek6 variants^87^, no potent Nek6 scaffolds have emerged. Perhaps the most promising Nek6 chemical starting point at present is an ATM inhibitor, compound 27g from St. Jude (Figure 11).^88^ This molecule inhibits ATM with an IC_50_ value of 1.2 µM, and was screened for kinase selectivity against 451 kinases at DiscoverX. Only 18 non-mutant kinases exhibited greater than 90% inhibition at a screening concentration of 3 µM. Among these off-targets were Nek6 and Nek9 with K_d_ values of 130 nM and 380 nM respectively. In this DiscoverX binding assay format, 27g demonstrated little to no binding to Nek1, Nek2, Nek3, Nek4, Nek5, and Nek11. The compound may be a weak inhibitor of Nek7 as well (78% inhibition at 3 µM), but no K_d_ value was reported. This quinazoline compound represents an interesting starting point to develop potent and selective Nek6, and perhaps Nek9, inhibitors.

**Figure 11.**
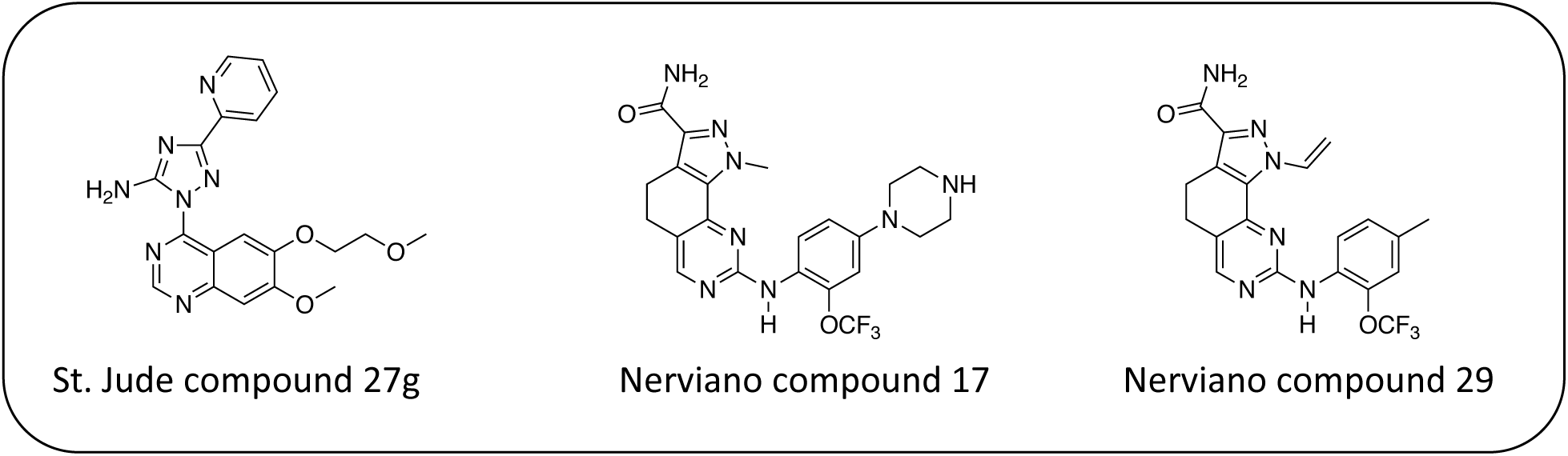
Nek6 inhibitors

Two compounds, Nerviano 29 and Nerviano 17 (Figure 11), designed for PLK1 demonstrate modest Nek6 inhibition.^89^ Nerviano17 had a Nek6 IC_50_ value of 595 nM, and only inhibited 4 kinases out of 43 screened with IC_50_ < 1 µM. Nerviano 29 was slightly more potent with Nek6 IC_50_ = 336 nM, and only inhibited 5 kinases out of the 43 screened with a potency under 1 µM. Like PLK1, Nek7 has a leucine at the hinge, which may explain why it tolerates the ortho-OCF3 group. Activity on the other Nek family members remains to be determined, and it is likely that only Nek7 and Nek10 would be candidates as targets because they too have a leucine in this position.

### Nek7

Initially identified as a regulator of the p70 ribosomal S6 kinase, Nek7 is required for microtubule nucleation, proper spindle assembly, and also plays a role in cytokinesis.^90–93^ Nek7 deficient mice displayed lethality in late embryogenesis or at early post-natal stages, and the mice displayed severe growth retardation, consistent with the role of this kinase in regulation of cell division.^94^ Nek7 activates the NLRP3 inflammasome and has thus been suggested as a novel target for the treatment of gout, atherosclerosis, Alzheimer’s disease, and Type II diabetes.^13^, ^95–97^ In addition, Nek7 was expressed in tumor samples from patients with breast cancer, colorectal cancer, cancer of the larynx, and melanoma,^85^ whereas it was not significantly expressed in matched normal tissue from these same patients.

There are no selective and potent small molecule inhibitors for Nek7 despite a deposited crystal structure (Table S2). The most promising compounds were originally synthesized as inhibitors of the JNK family of kinases. JNK-IN-1 (Figure 7) inhibited Nek7 activity by 80% inhibition at 10 µM (Table 10). Notably, this compound was only a modest inhibitor of Nek6 (44% control at 10 µM). Although both JNK-IN-2 and JNK-IN-7 inhibited Nek 7 by >90% of Nek7 activity, they have greater cross-Nek activity than JNK-IN-1, which showed potent inhibition only at Nek7. The structurally similar JNK-IN-8 lacks activity on Nek7, suggesting that the position of the electrophilic acrylamide group may be critical for Nek7 affinity. Both JNK-IN-2 and JNK-IN-7, which lack the methyl group of JNK-IN-1, showed promiscuous inhibition across the Nek family. Thus, the presence of methyl group in JNK-IN-1 may be required to achieve selectivity across the Nek family. Synthesis of additional analogues in this series would help elucidate SAR for Nek3, Nek5 and Nek7 and enable further biological studies to understand the consequence of inhibiting these targets.

### Nek8

First cloned and characterized in 2002, Nek8 is a tumor-associated kinase that is overexpressed in several human breast tumors.^98–99^ Nek8 also regulates the DNA replication fork protection protein RAD51 and is essential for the stability of replication fork and DNA repair.^100^ Mutations in Nek8 are associated with ciliary function disruptions and its dysregulation is linked to polycystic kidney disease and nephronophthisis.^101–102^

No compounds were identified as suitable starting points for the design of Nek8 inhibitors in our analysis. One possible reason could be that there are only 2 commercial screens available for Nek8,^4^ so it is not often included in reported kinase selectivity panels. Since Nek8 has a high level of active site homology with Nek5, Nek7 and Nek9 (Table 2), compounds that show affinity for these Nek family members, might also cross activity on Nek8. Screening of selected compounds belonging to chemotypes that have already shown binding affinity for other Nek family members may identify reasonable starting points for the synthesis of a selective Nek8 inhibitor.

### Nek9

Nek9 is involved in a mitotic cascade where it involving activates Nek6 and Nek7 during mitosis.^81^, ^103^ Nek9 is an enzymatic partner for the transcription factor FACT, that has a role in interphase progression.^104^ Nek9 has been shown to regulate microtubule nucleation, spindle organization, and cell cycle progression.^104–105^ Nek9 is also implicated in cancer as depletion of Nek9 in glioblastoma and kidney carcinoma cells resulted in impairment of cytokinesis and cell death.^106^ Gain of function mutations in Nek9 have been linked to the pathogenesis of *Nevus comedonicus* - a severe localized form of acne.^107^

GSK reported a series of imidazotriazines (Table 13) as PLK1 inhibitors.^108^ Several of these compounds are included in the PKIS and PKIS2 compounds sets and cross screening identified Nek9 as an off target. GW7731 showed robust Nek inhibition at both the 100 nM and 1000 nM screening concentrations. While replacement of the CF3 of GW7331 with the larger phenoxygroup in GW1893 rendered the compound inactive against Nek9, cross activity at Nek 6 emerged. UNC5391 is a member of PKIS2 with an indole in the place of the m-CF3 phenyl found in GW7331. Encouragingly, UNC5391 also shows Nek9 activity even though the results were obtained in a different assay format. Thus, based on these PKIS2 results, the compound may bind to several Nek family members. Given the narrow kinome profile of these three compounds, corroboration in two assay formats, and three sites for SAR exploration, further chemistry efforts to target other Nek family members are warranted.

**Table 13.**
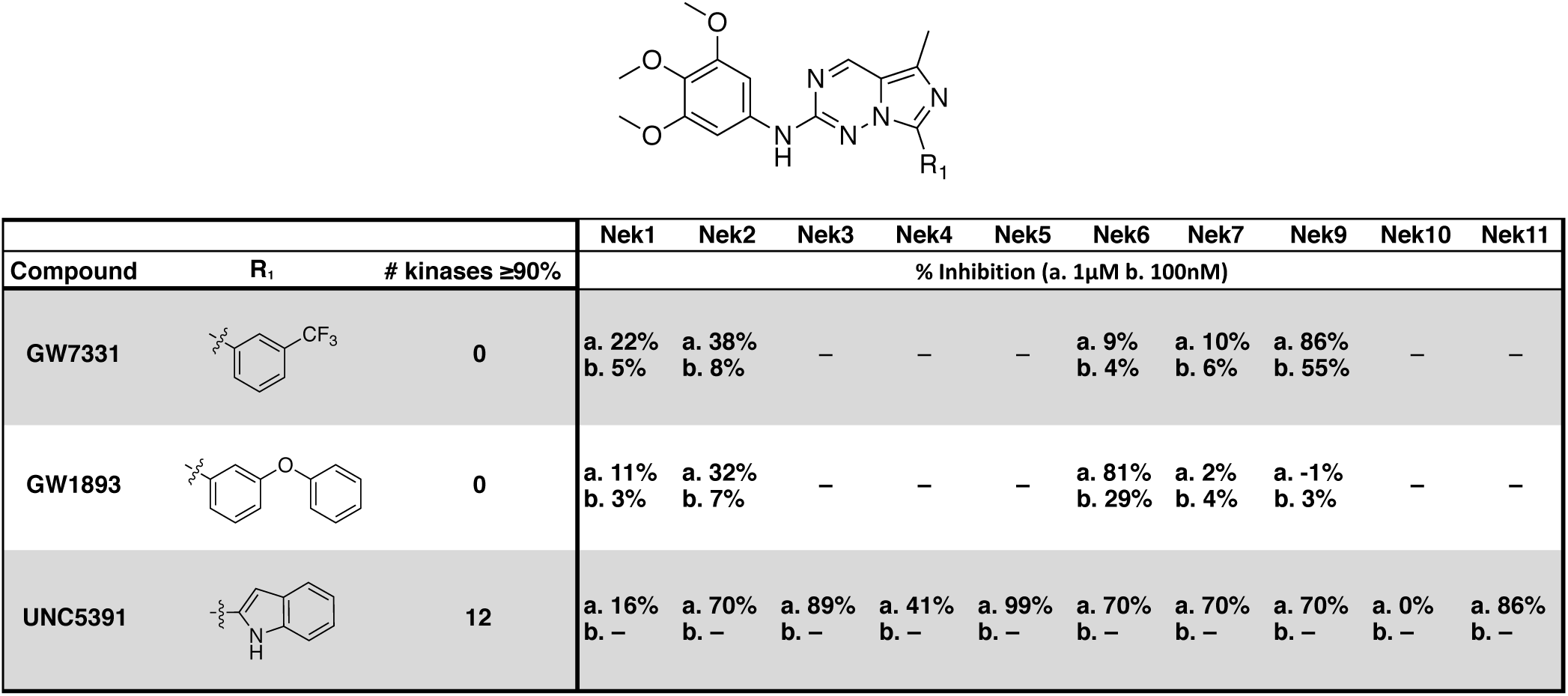
Imidazotriazine structures and percent inhibition values

### Nek10

Nek10 is one of the least studied Nek family members with a total of only 10 citations on PubMed (Table1). Despite this lack of attention, Nek10 has been identified as a potential causative gene for breast cancer through a Genome Wide Association Study (GWAS).^35^ Nek10 has also been shown to regulate the G2/M checkpoint in response to exposure of cells to UV irradiation and the DNA damaging agent etoposide.^109^ Importantly, Nek10 mutations have also been reported in several human cancers.^110^

Nek10 is the most divergent member of the Nek family with only 54% sequence identity to its nearest mammalian relatives (Nek6 and Nek7).^109^ Due to this difference we hypothesized that compounds inhibiting Nek10 would have minimal cross reactivity with other Nek family kinases. The Nek10 inhibition data from PKIS2 confirmed this prediction with only Nek3 inhibitors showing hints of cross activity (Figure 1b). One series that has Nek10 inhibition with little inhibition on the remainder of the Nek family was the cinnoline amides, (Figure 12a) originally published as BTK inhibitors.^111^ For BTK, the amide plays the role of hinge binding moiety. These compounds were contained in PKIS2, and broad kinase profiling uncovered the Nek10 activity (Figure 12b).^4^

**Figure 12.**
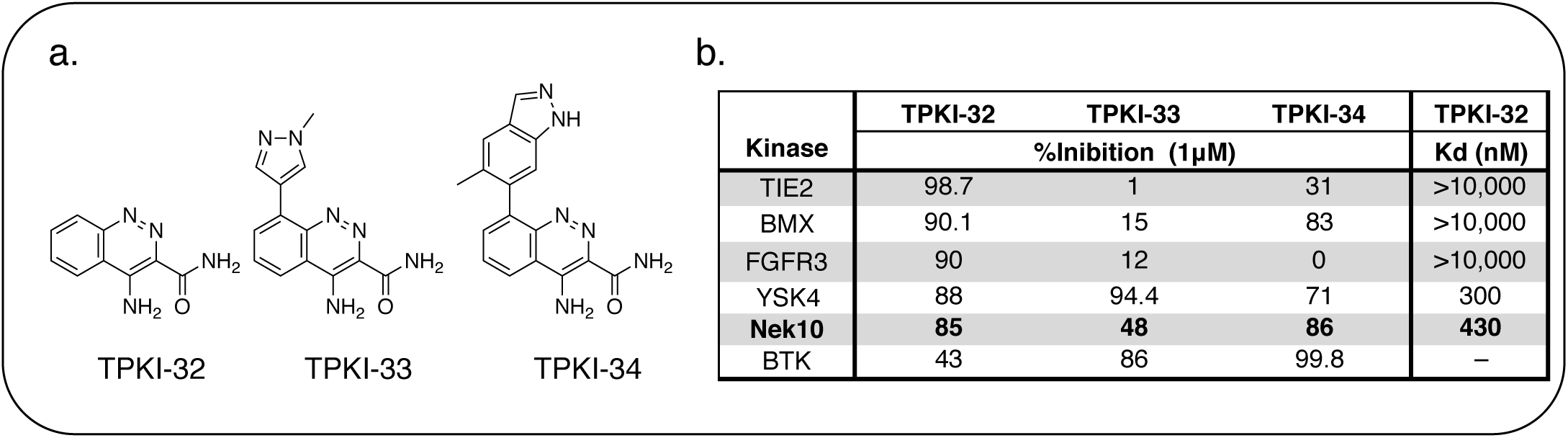
Takeda cinnoline amide a. compounds and b. Nek10 activity table

TPKI-32 binds to Nek10 with a K_d_ of 430nM and was also selective against the other Nek family kinases. Remarkably, TPKI-32 did not inhibit any other kinases in the large DiscoverX panel. TPKI-34, which incorporates a substituted indazole in the 8-position, retained activity on Nek10, but inhibited 21 kinases >90% inhibition. The increased promiscuity may arise from the introduction of the indazole as a second potential hinge binding group. The third member in the series, TPKI-33, has a methylated pyrazole in the 8-position. Nek10 binding decreased substantially and only three kinases were inhibited more than 90%. The very narrow kinase inhibition profile, promising potency, and evidence of SAR make this an exciting series for synthesis of selective Nek10 inhibitors.

Another potential Nek10 starting point, G1T28, a pyrrolopyrimidine designed as a CDK4/6 inhibitor (Figure 13) by G1 Therapeutics.

**Figure 13.**
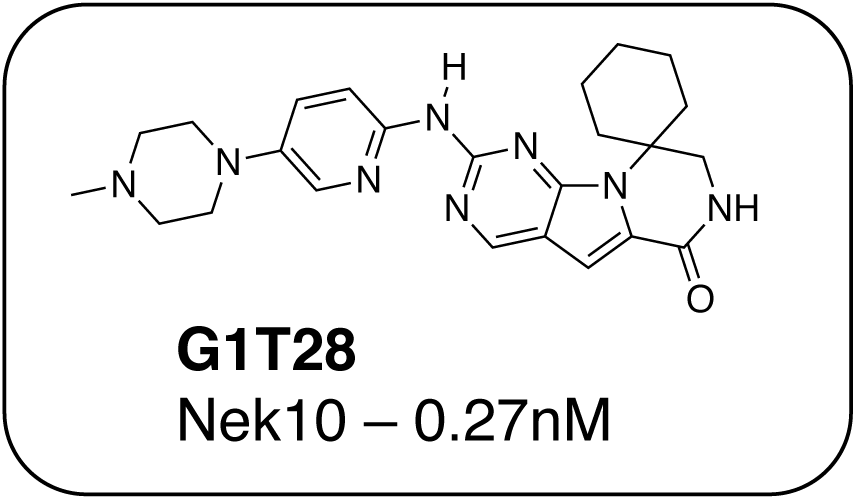
G1T28 (Trilaciclib) – a CDK4/6 inhibitor with Nek10 activity.

G1T28 has a fairly narrow selectivity profile with only 7% of the 468 membered DiscoverX KINOMEscan panel being inhibited >65% at a screening concentration of 100 nM. No other Nek family kinases exhibited greater than 90% inhibition. This compound inhibited Nek10 very potently (0.27 nM) in a biochemical assay. Broad exploration of this scaffold may lead to potent and selective Nek10 inhibitors with a sufficiently narrow profile to allow for exploration of Nek10 biology in cells.

3-anilino-4-arylmaleimides are another series of Nek10 inhibitors that were identified in the broad DiscoverX screening of PKIS2 (Table 14).^4^ These compounds were originally synthesized as GSK3B inhibitors^112^ and appear to be selective over the other Nek family members. Based on these results, the 3-COOH-4-Cl substituted aniline is a preferred group for Nek2 activity with a K_d_ of 460nM. A careful SAR study will be required to identify key features for Nek10 potency and kinome selectivity.

**Table 14.**
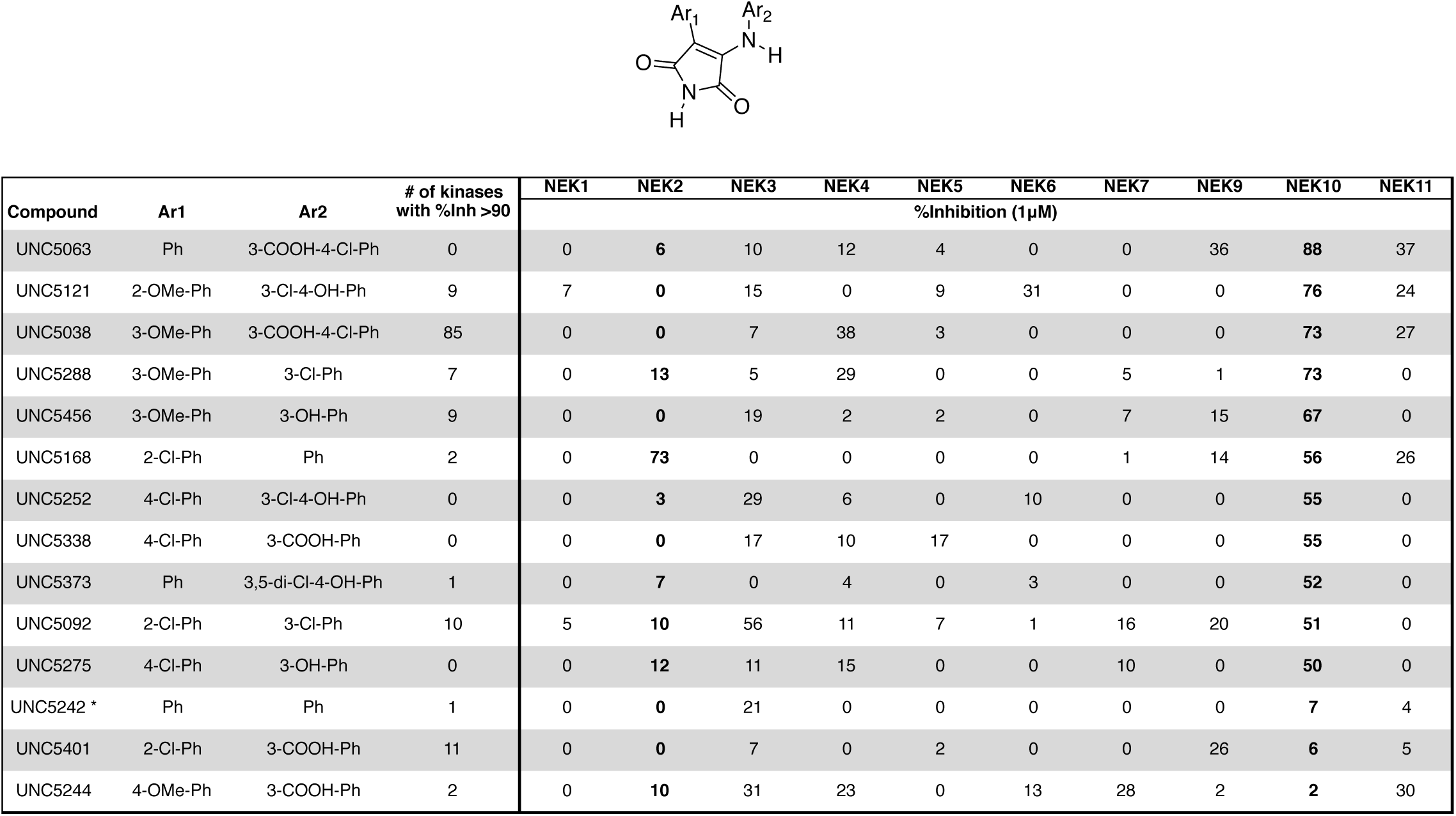
Nek10 SAR for 3-anilino-4-arylmaleimides from the Discoverx PKIS2 screen.

### Nek11

Nek11 is mainly implicated as a DNA damage/stress response kinase.^113–115^ Nek11 directly phosphorylates the checkpoint protein CDC25A and promotes its degradation.^116^ Inhibition of Nek11 results in cells entering cell cycle without a functional G1/S or G2/M checkpoint, which eventually leads to loss of genomic integrity and cell death.^117^ Furthermore, enhanced levels of Nek11 are found in HeLa cells following exposure to DNA damaging agents or replication inhibitors.^118^

While there are no reports of selective Nek11inhibitors, mining of kinase cross screening data uncovered a couple potential chemical starting points. PLX-4720^119^ (Figure 14) is a relatively selective BRAF inhibitor, with K_d_ values below 300 nM for only 3% of the kinases in a large panel including Nek11.^16^ PLX-4720 bound to Nek11 with a K_d_ value of 190 nM. PLX-4032 (Figure 14), also known as Vemurafenib or RG7204, is an approved BRAF inhibitor from the same azaindole scaffold.^120^ Interestingly, addition of the phenyl ring in the 5-position appeared to dampen binding to Nek11 (data from HMS LINCS dataset ID 20026). Dabrafenib (Figure 14), an FDA-approved BRAF inhibitor based on a different chemotype^121^, also bound to Nek11 (data from HMS LINCS ID 20131).

**Figure 14:**
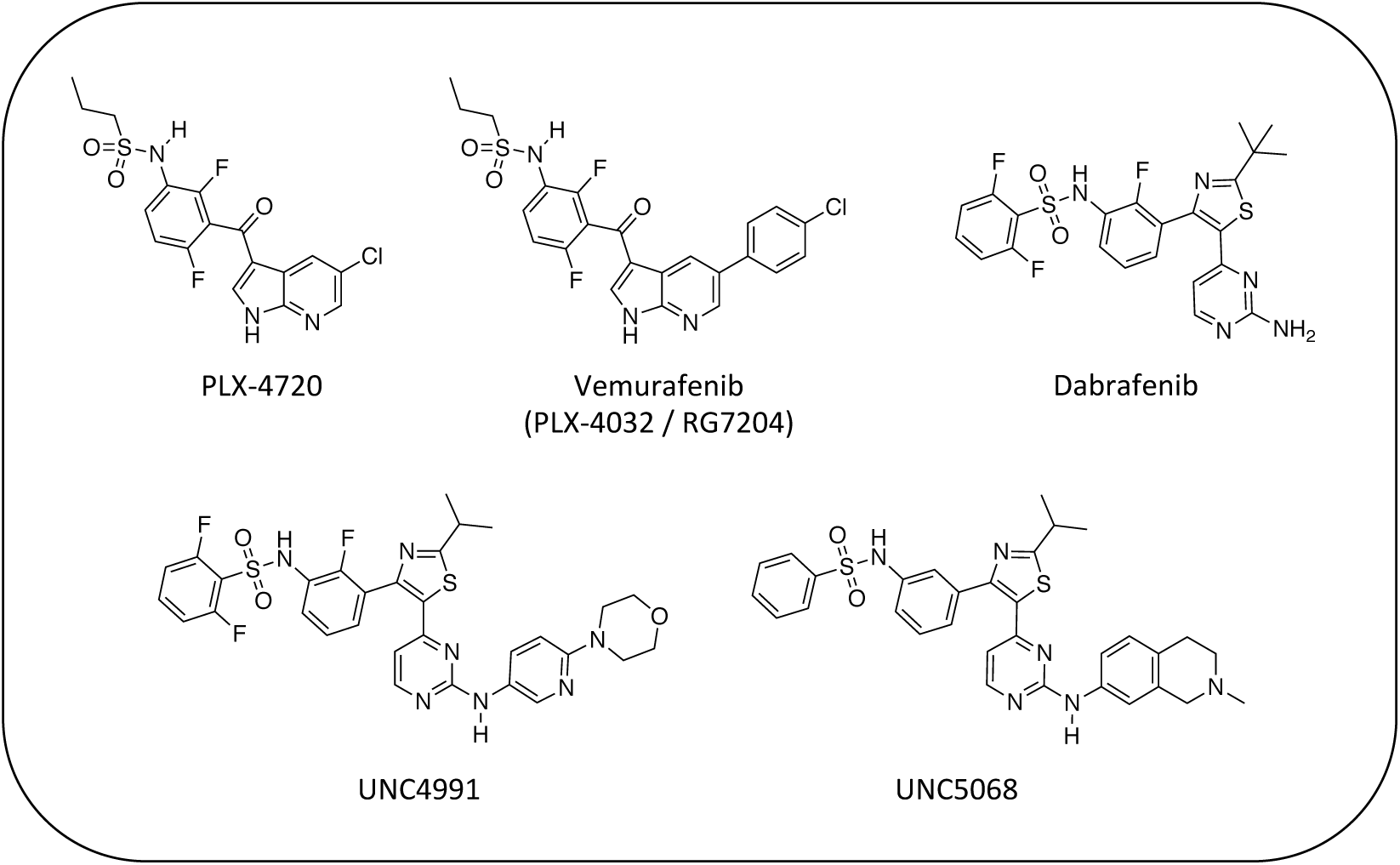
Structures of BRAF inhibitors and analogues for the design of Nek11 inhibitors

The reported data from single concentration screening at 10 µM against many members of the Nek family are shown in Table 15. Several other Nek family members are inhibited by dabrafenib, but require generation of K_d_ values to confirm binding. Further confirmation of the cross reactivity of the dabrafenib chemotype emerged from analysis of broad kinome screening data for the PKIS2. UNC4991 and UNC5068 both showed potent inhibition of Nek11 when screened at a 1 µM concentration. UNC4991 also inhibited several other Nek family members, suggesting that SAR may emerge if the series is further investigated.

**Table 15:** Nek family binding data for clinical BRAF inhibitors and analogues

| Table 15: Nek11 Inhibitors from BRAF series and related compounds |  |  |  |  |  |
| --- | --- | --- | --- | --- | --- |
| Compound | PLX-4720<sup>a</sup> | Vemurafenib<sup>b</sup> | Dabrafenib<sup>c</sup> | UNC4991<sup>d</sup> | UNC5068<sup>d</sup> |
| | % Control (10 $\mu\text{M}$) | | | % Control (1 $\mu\text{M}$) | |
| Nek1 | 72 | 100 | 14 | 18 | 86 |
| Nek2 | 49 (Kd = 5 $\mu\text{M}$) | 71 | 7 | 0.9 | 66 |
| Nek3 | 69 | 68 | 100 | 99 | 73 |
| Nek4 | 71 | 95 | 58 | 49 | 82 |
| Nek5 | 100 | 100 | 75 | 24 | 100 |
| Nek6 | 91 | 100 | 0.3 | 0.9 | 100 |
| Nek7 | 59 | 100 | 1.6 | 7.1 | 100 |
| Nek8 | - | - | - | - | - |
| Nek9 | 66 | 100 | 3.2 | 6.4 | 49 |
| Nek10 | - | - | 56 | 100 | 100 |
| Nek11 | 2 (Kd = 190 nM) | 30 | 0.2 | 1.4 | 2.3 |
| promiscuity | S(5%) at 10 $\mu\text{M}$ = 0.06 | S(5%) at 10 $\mu\text{M}$ = 0.07<br>S(300 nm) = 0.03<sup>e</sup> | S(5%) at 10 $\mu\text{M}$ = 0.19 | 137 kinases POC<10 | 110 kinases POC<10 |
| <sup>a</sup>values from HMS LINC dataset ID 20024 (single concentration and LINC HMS ID 20154 (Kd determinations))<br><sup>b</sup>values from HMS LINC dataset ID 20026<br><sup>c</sup>values from HMS LINC dataset ID 20131<br><sup>d</sup>values from Drewry et al., 2017<br><sup>e</sup>value from Davis et al., 2011 |  |  |  |  |  |

## Summary and Conclusions

Kinases are tractable targets, and investments in small molecule kinase inhibitors have led to increased understanding of kinase biology and therapies that benefit patients. In spite of this track record of success, much of the kinome is still understudied. Having a tool kit of kinase inhibitors that span the kinome will aid scientists in the identification of the right kinase target(s) for the right disease. To that end, we are working towards generation of such a tool set, the comprehensive kinase chemogenomic set (KCGS). Here we demonstrate that for the Nek family, a very poorly studied subset of kinases, careful and thorough data mining has allowed us to identify a broad range of potential chemical starting points and establish preliminary structure activity relationships. The ability to paint this landscape for kinases that have received scant attention is due to several factors, including:

a. the chemical connectivity of kinases – active site inhibitors for one kinase tend to cross-react with other kinases to varying degrees.
b. the widespread availability of ever growing broad kinase screening panels
c. the willingness of scientists to share these comprehensive data sets

We believe the analysis presented here and the identification of tractable chemical starting points will lead to the discovery of potent, selective, and cell active inhibitors for each Nek family member. These inhibitors will in turn lead to improved understanding of the intricacies of Nek biology, and the discovery of important medicines.

## Acknowledgements

This work was funded by the UNC Eshelman Institute for Innovation.

The SGC is a registered charity (number 1097737) that receives funds from AbbVie, Bayer Pharma AG, Boehringer Ingelheim, Canada Foundation for Innovation, Eshelman Institute for Innovation, Genome Canada, Innovative Medicines Initiative (EU/EFPIA) [ULTRA-DD grant no. 115766], Janssen, Merck & Co., Novartis Pharma AG, Ontario Ministry of Economic Development and Innovation, Pfizer, São Paulo Research Foundation-FAPESP, Takeda, and Wellcome Trust [092809/Z/10/Z].

The authors thank Tim Willson (SGC) for helpful discussions and feedback.

GlaxoSmithKline, Pfizer, and Takeda are gratefully acknowledged for donation of PKIS2 compounds.

## Supplementary Information Available

Table S1 – Kinase active site residue sequence alignment for the Nek family of kinases

Figure S1 – Kinase selectivity bar chart for PKIS2 data for DiscoverX screening data at 1 µM

Table S2 – Nek family crystal structure availability

## References

1. Rask-Andersen, M.; Zhang, J.; Fabbro, D.; Schioeth, H. B., Advances in kinase targeting: current clinical use and clinical trials. Trends Pharmacol. Sci. 2014, 35 (11), 604–620.

2. Wu, P.; Nielsen, T. E.; Clausen, M. H., FDA-approved small-molecule kinase inhibitors. Trends Pharmacol. Sci. 2015, 36 (7), 422–439.

3. Fedorov, O.; Mueller, S.; Knapp, S., The (un)targeted cancer kinome. Nat. Chem. Biol. 2010, 6 (3), 166–169.

4. Drewry, D. H.; Wells, C. I.; Andrews, D. M.; Angell, R.; Al-Ali, H.; Axtman, A. D.; Capuzzi, S. J.; Elkins, J. M.; Ettmayer, P.; Frederiksen, M.; Gileadi, O.; Gray, N.; Hooper, A.; Knapp, S.; Laufer, S.; Luecking, U.; Muller, S.; Muratov, E.; Denny, R. A.; Saikatendu, K. S.; Treiber, D. K.; Zuercher, W. J.; Willson, T. M., Progress Towards a Public Chemogenomic Set for Protein Kinases and a Call for Contributions. bioRxiv 2017.

5. Yu, D.; Hung, M. C., Overexpression of ErbB2 in cancer and ErbB2-targeting strategies. Oncogene 2000, 19 (53), 6115–21.

6. Oakley, B. R.; Morris, N. R., A mutation in Aspergillus nidulans that blocks the transition from interphase to prophase. J Cell Biol 1983, 96 (4), 1155–8.

7. Morris, N. R., Mitotic mutants of Aspergillus nidulans. Genet Res 1975, 26 (3), 237–54.

8. Quarmby, L. M.; Mahjoub, M. R., Caught Nek-ing: cilia and centrioles. J. Cell Sci. 2005, 118 (22), 5161–5169.

9. Fry, A. M.; O'Regan, L.; Sabir, S. R.; Bayliss, R., Cell cycle regulation by the NEK family of protein kinases. J. Cell Sci. 2012, 125 (19), 4423–4433.

10. Bamborough, P.; Drewry, D.; Harper, G.; Smith, G. K.; Schneider, K., Assessment of Chemical Coverage of Kinome Space and Its Implications for Kinase Drug Discovery. Journal of Medicinal Chemistry 2008, 51 (24), 7898–7914.

11. Posy, S. L.; Hermsmeier, M. A.; Vaccaro, W.; Ott, K. H.; Todderud, G.; Lippy, J. S.; Trainor, G. L.; Loughney, D. A.; Johnson, S. R., Trends in kinase selectivity: insights for target class-focused library screening. J Med Chem 2011, 54 (1), 54–66.

12. Gao, Y.; Davies, S. P.; Augustin, M.; Woodward, A.; Patel, U. A.; Kovelman, R.; Harvey, K. J., A broad activity screen in support of a chemogenomic map for kinase signalling research and drug discovery. Biochem J 2013, 451 (2), 313–28.

13. Elkins, J. M.; Fedele, V.; Szklarz, M.; Abdul Azeez, K. R.; Salah, E.; Mikolajczyk, J.; Romanov, S.; Sepetov, N.; Huang, X.-P.; Roth, B. L.; Al Haj Zen, A.; Fourches, D.; Muratov, E.; Tropsha, A.; Morris, J.; Teicher, B. A.; Kunkel, M.; Polley, E.; Lackey, K. E.; Atkinson, F. L.; Overington, J. P.; Bamborough, P.; Muller, S.; Price, D. J.; Willson, T. M.; Drewry, D. H.; Knapp, S.; Zuercher, W. J., Comprehensive characterization of the Published Kinase Inhibitor Set. Nat Biotech 2016, 34 (1), 95–103.

14. Anastassiadis, T.; Deacon, S. W.; Devarajan, K.; Ma, H.; Peterson, J. R., Comprehensive assay of kinase catalytic activity reveals features of kinase inhibitor selectivity. Nat Biotechnol 2011, 29 (11), 1039–45.

15. Bamborough, P.; Drewry, D.; Harper, G.; Smith, G. K.; Schneider, K., Assessment of chemical coverage of kinome space and its implications for kinase drug discovery. J Med Chem 2008, 51 (24), 7898–914.

16. Davis, M. I.; Hunt, J. P.; Herrgard, S.; Ciceri, P.; Wodicka, L. M.; Pallares, G.; Hocker, M.; Treiber, D. K.; Zarrinkar, P. P., Comprehensive analysis of kinase inhibitor selectivity. Nat Biotechnol 2011, 29 (11), 1046–51.

17. Braselmann, S.; Taylor, V.; Zhao, H.; Wang, S.; Sylvain, C.; Baluom, M.; Qu, K.; Herlaar, E.; Lau, A.; Young, C.; Wong, B. R.; Lovell, S.; Sun, T.; Park, G.; Argade, A.; Jurcevic, S.; Pine, P.; Singh, R.; Grossbard, E. B.; Payan, D. G.; Masuda, E. S., R406, an orally available spleen tyrosine kinase inhibitor blocks fc receptor signaling and reduces immune complex-mediated inflammation. J Pharmacol Exp Ther 2006, 319 (3), 998–1008.

18. Pratz, K. W.; Cortes, J.; Roboz, G. J.; Rao, N.; Arowojolu, O.; Stine, A.; Shiotsu, Y.; Shudo, A.; Akinaga, S.; Small, D.; Karp, J. E.; Levis, M., A pharmacodynamic study of the FLT3 inhibitor KW-2449 yields insight into the basis for clinical response. Blood 2009, 113 (17), 3938–3946.

19. Zhao, Z.; Wu, H.; Wang, L.; Liu, Y.; Knapp, S.; Liu, Q.; Gray, N. S., Exploration of type II binding mode: A privileged approach for kinase inhibitor focused drug discovery? ACS Chem Biol 2014, 9 (6), 1230–41.

20. Tyner, J. W.; Bumm, T. G.; Deininger, J.; Wood, L.; Aichberger, K. J.; Loriaux, M. M.; Druker, B. J.; Burns, C. J.; Fantino, E.; Deininger, M. W., CYT387, a novel JAK2 inhibitor, induces hematologic responses and normalizes inflammatory cytokines in murine myeloproliferative neoplasms. Blood 2010, 115 (25), 5232.

21. Höfener, M.; Pachl, F.; Kuster, B.; Sewald, N., Inhibitor-based affinity probes for the investigation of JAK signaling pathways. PROTEOMICS 2015, 15 (17), 3066–3074.

22. Liang, Y.; Meng, D.; Zhu, B.; Pan, J., Mechanism of ciliary disassembly. Cell Mol Life Sci 2016, 73 (9), 1787–802.

23. Shalom, O.; Shalva, N.; Altschuler, Y.; Motro, B., The mammalian Nek1 kinase is involved in primary cilium formation. FEBS Lett. 2008, 582 (10), 1465–1470.

24. Upadhya, P.; Birkenmeier, E. H.; Birkenmeier, C. S.; Barker, J. E., Mutations in a NIMA-related kinase gene, Nek1, cause pleiotropic effects including a progressive polycystic kidney disease in mice. Proc. Natl. Acad. Sci. U. S. A. 2000, 97 (1), 217–221.

25. Mahjoub, M. R.; Trapp, M. L.; Quarmby, L. M., NIMA-related kinases defective in murine models of polycystic kidney diseases localize to primary cilia and centrosomes. J. Am. Soc. Nephrol. 2005, 16 (12), 3485–3489.

26. White, M. C.; Quarmby, L. M., The NIMA-family kinase, Nek1 affects the stability of centrosomes and ciliogenesis. BMC Cell Biol. 2008, 9, No pp. given.

27. Thiel, C.; Kessler, K.; Giessl, A.; Dimmler, A.; Shalev, S. A.; von der Haar, S.; Zenker, M.; Zahnleiter, D.; Stoess, H.; Beinder, E.; Abou Jamra, R.; Ekici, A. B.; Schroeder-Kress, N.; Aigner, T.; Kirchner, T.; Reis, A.; Brandstaetter, J. H.; Rauch, A., NEK1 Mutations Cause Short-Rib Polydactyly Syndrome Type Majewski. Am. J. Hum. Genet. 2010, 88 (1), 106–114.

28. Chen, C.-P.; Chang, T.-Y.; Chen, C.-Y.; Wang, T.-Y.; Tsai, F.-J.; Wu, P.-C.; Chern, S.-R.; Wang, W., Short rib-polydactyly syndrome type II (Majewski): prenatal diagnosis, perinatal imaging findings and molecular analysis of the NEK1 gene. Taiwan J Obstet Gynecol 2012, 51 (1), 100–5.

29. El Hokayem, J.; Huber, C.; Couve, A.; Aziza, J.; Baujat, G.; Bouvier, R.; Cavalcanti, D. P.; Collins, F. A.; Cordier, M.-P.; Delezoide, A.-L.; Gonzales, M.; Johnson, D.; Le Merrer, M.; Levy-Mozziconacci, A.; Loget, P.; Martin-Coignard, D.; Martinovic, J.; Mortier, G. R.; Perez, M.-J.; Roume, J.; Scarano, G.; Munnich, A.; Cormier-Daire, V., NEK1 and DYNC2H1 are both involved in short rib polydactyly Majewski type but not in Beemer Langer cases. J. Med. Genet. 2012, 49 (4), 227–233.

30. Zhu, J.; Cai, Y.; Liu, P.; Zhao, W., Frequent Nek1 overexpression in human gliomas. Biochem. Biophys. Res. Commun. 2016, 476 (4), 522–527.

31. Polci, R.; Peng, A.; Chen, P.-L.; Riley, D. J.; Chen, Y., NIMA-Related Protein Kinase 1 Is Involved Early in the Ionizing Radiation-Induced DNA Damage Response. Cancer Res. 2004, 64 (24), 8800–8803.

32. Chen, Y.; Chen, P.-L.; Chen, C.-F.; Jiong, X.; Riley, D. J., Never-in-mitosis related kinase 1 functions in DNA damage response and checkpoint control. Cell Cycle 2008, 7 (20), 3194–3201.

33. Pelegrini, A. L.; Moura, D. J.; Brenner, B. L.; Ledur, P. F.; Maques, G. P.; Henriques, J. A. P.; Saffi, J.; Lenz, G., Nek1 silencing slows down DNA repair and blocks DNA damage-induced cell cycle arrest. Mutagenesis 2010, 25 (5), 447–454.

34. Yamamoto, N., The Orally Available Spleen Tyrosine Kinase Inhibitor 2-[7-(3,4-Dimethoxyphenyl)-imidazo[1,2-c]pyrimidin-5-ylamino]-nicotinamide Dihydrochloride (BAPY 61-3606) Blocks Antigen-Induced Airway Inflmmation in Rodents. The Journal of Pharmacology and Experimental Therapeutics 2003, 306 (3), 1174–1181.

35. Ahmed, S.; Thomas, G.; Ghoussaini, M.; Healey, C. S.; Humphreys, M. K.; Platte, R.; Morrison, J.; Maranian, M.; Pooley, K. A.; Luben, R.; Eccles, D.; Evans, D. G.; Fletcher, O.; Johnson, N.; dos Santos Silva, I.; Peto, J.; Stratton, M. R.; Rahman, N.; Jacobs, K.; Prentice, R.; Anderson, G. L.; Rajkovic, A.; Curb, J. D.; Ziegler, R. G.; Berg, C. D.; Buys, S. S.; McCarty, C. A.; Feigelson, H. S.; Calle, E. E.; Thun, M. J.; Diver, W. R.; Bojesen, S.; Nordestgaard, B. G.; Flyger, H.; Doerk, T.; Schuermann, P.; Hillemanns, P.; Karstens, J. H.; Bogdanova, N. V.; Antonenkova, N. N.; Zalutsky, I. V.; Bermisheva, M.; Fedorova, S.; Khusnutdinova, E.; Search; Kang, D.; Yoo, K.-Y.; Noh, D. Y.; Ahn, S.-H.; Devilee, P.; van Asperen, C. J.; Tollenaar, R. A. E. M.; Seynaeve, C.; Garcia-Closas, M.; Lissowska, J.; Brinton, L.; Peplonska, B.; Nevanlinna, H.; Heikkinen, T.; Aittomaeki, K.; Blomqvist, C.; Hopper, J. L.; Southey, M. C.; Smith, L.; Spurdle, A. B.; Schmidt, M. K.; Broeks, A.; van Hien, R. R.; Cornelissen, S.; Milne, R. L.; Ribas, G.; Gonzalez-Neira, A.; Benitez, J.; Schmutzler, R. K.; Burwinkel, B.; Bartram, C. R.; Meindl, A.; Brauch, H.; Justenhoven, C.; Hamann, U.; Chang-Claude, J.; Hein, R.; Wang-Gohrke, S.; Lindblom, A.; Margolin, S.; Mannermaa, A.; Kosma, V.-M.; Kataja, V.; Olson, J. E.; Wang, X.; Fredericksen, Z.; Giles, G. G.; Severi, G.; Baglietto, L.; English, D. R.; Hankinson, S. E.; Cox, D. G.; Kraft, P.; Vatten, L. J.; Hveem, K.; Kumle, M.; Sigurdson, A.; Doody, M.; Bhatti, P.; Alexander, B. H.; Hooning, M. J.; van den Ouweland, A. M. W.; Oldenburg, R. A.; Schutte, M.; Hall, P.; Czene, K.; Liu, J.; Li, Y.; Cox, A.; Elliott, G.; Brock, I.; Reed, M. W. R.; Shen, C.-Y.; Yu, J.-C.; Hsu, G.-C.; Chen, S.-T.; Anton-Culver, H.; Ziogas, A.; Andrulis, I. L.; Knight, J. A.; Beesley, J.; Goode, E. L.; Couch, F.; Chenevix-Trench, G.; Hoover, R. N.; Ponder, B. A. J.; Hunter, D. J.; Pharoah, P. D. P.; Dunning, A. M.; Chanock, S. J.; Easton, D. F., Newly discovered breast cancer susceptibility loci on 3p24 and 17q23.2. Nat. Genet. 2009, 41 (5), 585–590.

36. Metz, J. T.; Johnson, E. F.; Soni, N. B.; Merta, P. J.; Kifle, L.; Hajduk, P. J., Navigating the kinome. Nat Chem Biol 2011, 7 (4), 200–202.

37. Lau, K. S.; Zhang, T.; Kendall, K. R.; Lauffenburger, D.; Gray, N. S.; Haigis, K. M., BAY61-3606 affects the viability of colon cancer cells in a genotype-directed manner. PLoS One 2012, 7 (7), e41343.

38. Fry, A. M.; Meraldi, P.; Nigg, E. A., A centrosomal function for the human Nek2 protein kinase, a member of the NIMA family of cell cycle regulators. EMBO J. 1998, 17 (2), 470–481.

39. Fry, A. M.; Mayor, T.; Meraldi, P.; Stierhof, Y.-D.; Tanaka, K.; Nigg, E. A., C-Nap1, a novel centrosomal coiled-coil protein and candidate substrate of the cell cycle-regulated protein kinase Nek2. J. Cell Biol. 1998, 141 (7), 1563–1574.

40. Helps, N. R.; Luo, X.; Barker, H. M.; Cohen, P. T. W., NIMA-related kinase 2 (Nek2), a cell-cycle-regulated protein kinase localized to centrosomes, is complexed to protein phosphatase 1. Biochem. J. 2000, 349 (2), 509–518.

41. Bahe, S.; Stierhof, Y.-D.; Wilkinson, C. J.; Leiss, F.; Nigg, E. A., Rootletin forms centriole-associated filaments and functions in centrosome cohesion. J. Cell Biol. 2005, 171 (1), 27–33.

42. Rapley, J.; Baxter, J. E.; Blot, J.; Wattam, S. L.; Casenghi, M.; Meraldi, P.; Nigg, E. A.; Fry, A. M., Coordinate regulation of the mother centriole component Nlp by Nek2 and Plk1 protein kinases. Mol. Cell. Biol. 2005, 25 (4), 1309–1324.

43. Lou, Y.; Yao, J.; Zereshki, A.; Dou, Z.; Ahmed, K.; Wang, H.; Hu, J.; Wang, Y.; Yao, X., NEK2A Interacts with MAD1 and Possibly Functions as a Novel Integrator of the Spindle Checkpoint Signaling. J. Biol. Chem. 2004, 279 (19), 20049–20057.

44. Chen, Y.; Riley, D. J.; Zheng, L.; Chen, P.-L.; Lee, W.-H., Phosphorylation of the Mitotic Regulator Protein Hec1 by Nek2 Kinase is Essential for Faithful Chromosome Segregation. J. Biol. Chem. 2002, 277 (51), 49408–49416.

45. Tsunoda, N.; Kokuryo, T.; Oda, K.; Senga, T.; Yokoyama, Y.; Nagino, M.; Nimura, Y.; Hamaguchi, M., Nek2 as a novel molecular target for the treatment of breast carcinoma. Cancer Sci. 2009, 100 (1), 111–116.

46. Wu, S.-M.; Lin, S.-L.; Lee, K.-Y.; Chuang, H.-C.; Feng, P.-H.; Cheng, W.-L.; Liao, C.-J.; Chi, H.-C.; Lin, Y.-H.; Tsai, C.-Y.; Chen, W.-J.; Yeh, C.-T.; Lin, K.-H., Hepatoma cell functions modulated by NEK2 are associated with liver cancer progression. Int. J. Cancer 2017, 140 (7), 1581–1596.

47. Lu, L.; Zhai, X.; Yuan, R., Clinical significance and prognostic value of Nek2 protein expression in colon cancer. Int J Clin Exp Pathol 2015, 8 (11), 15467–73.

48. Xia, J.; Machin, R. F.; Gu, Z.; Zhan, F., Role of NEK2A in human cancer and its therapeutic potentials. BioMed Res. Int. 2015, 1–13.

49. Emmitte, K. A.; Adjebang, G. M.; Andrews, C. W.; Alberti, J. G. B.; Bambal, R.; Chamberlain, S. D.; Davis-Ward, R. G.; Dickson, H. D.; Hassler, D. F.; Hornberger, K. R.; Jackson, J. R.; Kuntz, K. W.; Lansing, T. J.; Mook Jr, R. A.; Nailor, K. E.; Pobanz, M. A.; Smith, S. C.; Sung, C.-M.; Cheung, M., Design of potent thiophene inhibitors of polo-like kinase 1 with improved solubility and reduced protein binding. Bioorganic & Medicinal Chemistry Letters 2009, 19 (6), 1694–1697.

50. Solanki, S.; Innocenti, P.; Mas-Droux, C.; Boxall, K.; Barillari, C.; van Montfort, R. L. M.; Aherne, G. W.; Bayliss, R.; Hoelder, S., Benzimidazole Inhibitors Induce a DFG-Out Conformation of Never in Mitosis Gene A-Related Kinase 2 (Nek2) without Binding to the Back Pocket and Reveal a Nonlinear Structure-Activity Relationship. Journal of Medicinal Chemistry 2011, 54 (6), 1626–1639.

51. Xi, J.-B.; Fang, Y.-F.; Frett, B.; Zhu, M.-L.; Zhu, T.; Kong, Y.-N.; Guan, F.-J.; Zhao, Y.; Zhang, X.-W.; Li, H.-y.; Ma, M.-L.; Hu, W., Structure-based design and synthesis of imidazo[1,2-a]pyridine derivatives as novel and potent Nek2 inhibitors with in vitro and in vivo antitumor activities. European Journal of Medicinal Chemistry 2017, 126, 1083-1106.

52. Whelligan, D. K.; Solanki, S.; Taylor, D.; Thomson, D. W.; Cheung, K.-M. J.; Boxall, K.; Mas-Droux, C.; Barillari, C.; Burns, S.; Grummitt, C. G.; Collins, I.; van Montfort, R. L. M.; Aherne, G. W.; Bayliss, R.; Hoelder, S., Aminopyrazine Inhibitors Binding to an Unusual Inactive Conformation of the Mitotic Kinase Nek2: SAR and Structural Characterization. Journal of Medicinal Chemistry 2010, 53 (21), 7682–7698.

53. Innocenti, P.; Cheung, K.-M. J.; Solanki, S.; Mas-Droux, C.; Rowan, F.; Yeoh, S.; Boxall, K.; Westlake, M.; Pickard, L.; Hardy, T.; Baxter, J. E.; Aherne, G. W.; Bayliss, R.; Fry, A. M.; Hoelder, S., Design of Potent and Selective Hybrid Inhibitors of the Mitotic Kinase Nek2: Structure–Activity Relationship, Structural Biology, and Cellular Activity. Journal of Medicinal Chemistry 2012, 55 (7), 3228–3241.

54. Henise, J. C.; Taunton, J., Irreversible Nek2 Kinase Inhibitors with Cellular Activity. Journal of Medicinal Chemistry 2011, 54 (12), 4133–4146.

55. Coxon, C. R.; Wong, C.; Bayliss, R.; Boxall, K.; Carr, K. H.; Fry, A. M.; Hardcastle, I. R.; Matheson, C. J.; Newell, D. R.; Sivaprakasam, M.; Thomas, H.; Turner, D.; Yeoh, S.; Wang, L. Z.; Griffin, R. J.; Golding, B. T.; Cano, C., Structure-guided design of purine-based probes for selective Nek2 inhibition. Oncotarget 2017, 8 (12), 19089–19124.

56. Meng, L.; Carpenter, K.; Mollard, A.; Vankayalapati, H.; Warner, S. L.; Sharma, S.; Tricot, G.; Zhan, F.; Bearss, D. J., Inhibition of Nek2 by Small Molecules Affects Proteasome Activity. BioMed Research International 2014, 2014, 13.

57. Kothe, M.; Kohls, D.; Low, S.; Coli, R.; Rennie, G. R.; Feru, F.; Kuhn, C.; Ding, Y.-H., Research Article: Selectivity-determining Residues in Plk1. Chemical Biology & Drug Design 2007, 70 (6), 540–546.

58. Rabindran, S. K.; Discafani, C. M.; Rosfjord, E. C.; Baxter, M.; Floyd, M. B.; Golas, J.; Hallett, W. A.; Johnson, B. D.; Nilakantan, R.; Overbeek, E.; Reich, M. F.; Shen, R.; Shi, X.; Tsou, H.-R.; Wang, Y.-F.; Wissner, A., Antitumor Activity of HKI-272, an Orally Active, Irreversible Inhibitor of the HER-2 Tyrosine Kinase. Cancer Research 2004, 64 (11), 3958.

59. Das, T. K.; Dana, D.; Paroly, S. S.; Perumal, S. K.; Singh, S.; Jhun, H.; Pendse, J.; Cagan, R. L.; Talele, T. T.; Kumar, S., Centrosomal kinase Nek2 cooperates with oncogenic pathways to promote metastasis. Oncogenesis 2013, 2, e69.

60. Fidanze, S. D.; Erickson, S. A.; Wang, G. T.; Mantei, R.; Clark, R. F.; Sorensen, B. K.; Bamaung, N. Y.; Kovar, P.; Johnson, E. F.; Swinger, K. K.; Stewart, K. D.; Zhang, Q.; Tucker, L. A.; Pappano, W. N.; Wilsbacher, J. L.; Wang, J.; Sheppard, G. S.; Bell, R. L.; Davidsen, S. K.; Hubbard, R. D., Imidazo[2,1-b]thiazoles: Multitargeted inhibitors of both the insulin-like growth factor receptor and members of the epidermal growth factor family of receptor tyrosine kinases. Bioorganic & Medicinal Chemistry Letters 2010, 20 (8), 2452–2455.

61. Axten, J. M.; Medina, J. R.; Feng, Y.; Shu, A.; Romeril, S. P.; Grant, S. W.; Li, W. H. H.; Heerding, D. A.; Minthorn, E.; Mencken, T.; Atkins, C.; Liu, Q.; Rabindran, S.; Kumar, R.; Hong, X.; Goetz, A.; Stanley, T.; Taylor, J. D.; Sigethy, S. D.; Tomberlin, G. H.; Hassell, A. M.; Kahler, K. M.; Shewchuk, L. M.; Gampe, R. T., Discovery of 7-Methyl-5-(1-{[3-(trifluoromethyl)phenyl]acetyl}-2,3-dihydro-1H-indol-5-yl)-7H-pyrrolo[2,3-d]pyrimidin-4-amine (GSK2606414), a Potent and Selective First-in-Class Inhibitor of Protein Kinase R (PKR)-like Endoplasmic Reticulum Kinase (PERK). Journal of Medicinal Chemistry 2012, 55 (16), 7193–7207.

62. Tanaka, K.; Nigg, E. A., Cloning and characterization of the murine Nek3 protein kinase, a novel member of the NIMA family of putative cell cycle regulators. J. Biol. Chem. 1999, 274 (19), 13491–7.

63. Miller, S. L.; DeMaria, J. E.; Freier, D. O.; Riegel, A. M.; Clevenger, C. V., Novel association of Vav2 and Nek3 modulates signaling through the human prolactin receptor. Molecular endocrinology (Baltimore, Md.) 2005, 19 (4), 939–49.

64. Miller, S. L.; Antico, G.; Raghunath, P. N.; Tomaszewski, J. E.; Clevenger, C. V., Nek3 kinase regulates prolactin-mediated cytoskeletal reorganization and motility of breast cancer cells. Oncogene 2007, 26 (32), 4668–4678.

65. Harrington, K. M.; Clevenger, C. V., Identification of NEK3 Kinase Threonine 165 as a Novel Regulatory Phosphorylation Site That Modulates Focal Adhesion Remodeling Necessary for Breast Cancer Cell Migration. J. Biol. Chem. 2016, 291 (41), 21388–21406.

66. Chang, J.; Baloh, R. H.; Milbrandt, J., The NIMA-family kinase Nek3 regulates microtubule acetylation in neurons. J. Cell Sci. 2009, 122 (13), 2274–2282.

67. Bakre, A.; Andersen, L. E.; Meliopoulos, V.; Coleman, K.; Yan, X.; Brooks, P.; Crabtree, J.; Tompkins, S. M.; Tripp, R. A., Identification of host kinase genes required for influenza virus replication and the regulatory role of microRNAs. PLoS One 2013, 8 (6), e66796.

68. Zhang, T.; Inesta-Vaquera, F.; Niepel, M.; Zhang, J.; Ficarro, S. B.; Machleidt, T.; Xie, T.; Marto, J. A.; Kim, N.; Sim, T.; Laughlin, J. D.; Park, H.; LoGrasso, P. V.; Patricelli, M.; Nomanbhoy, T. K.; Sorger, P. K.; Alessi, D. R.; Gray, N. S., Discovery of potent and selective covalent inhibitors of JNK. Chem Biol 2012, 19 (1), 140–54.

69. Kamenecka, T.; Jiang, R.; Song, X.; Duckett, D.; Chen, W.; Ling, Y. Y.; Habel, J.; Laughlin, J. D.; Chambers, J.; Figuera-Losada, M.; Cameron, M. D.; Lin, L.; Ruiz, C. H.; LoGrasso, P. V., Synthesis, biological evaluation, X-ray structure, and pharmacokinetics of aminopyrimidine c-jun-N-terminal kinase (JNK) inhibitors. J Med Chem 2010, 53 (1), 419–31.

70. Doles, J.; Hemann, M. T., Nek4 Status Differentially Alters Sensitivity to Distinct Microtubule Poisons. Cancer Res. 2010, 70 (3), 1033–1041.

71. Nguyen, C. L.; Possemato, R.; Bauerlein, E. L.; Xie, A.; Scully, R.; Hahn, W. C., Nek4 regulates entry into replicative senescence and the response to DNA damage in human fibroblasts. Mol. Cell. Biol. 2012, 32 (19), 3963–3977.

72. Park, S. J.; Jo, D. S.; Jo, S. Y.; Shin, D. W.; Shim, S.; Jo, Y. K.; Shin, J. H.; Ha, Y. J.; Jeong, S. Y.; Hwang, J. J.; Kim, Y. S.; Suh, Y. A.; Chang, J. W.; Kim, J. C.; Cho, D. H., Inhibition of never in mitosis A (NIMA)-related kinase-4 reduces survivin expression and sensitizes cancer cells to TRAIL-induced cell death. Oncotarget 2016, 7 (40), 65957–65967.

73. Pastrana-Mena, R.; Mathias, D. K.; Delves, M.; Rajaram, K.; King, J. G.; Yee, R.; Trucchi, B.; Verotta, L.; Dinglasan, R. R., A Malaria Transmission-Blocking (+)-Usnic Acid Derivative Prevents Plasmodium Zygote-to-Ookinete Maturation in the Mosquito Midgut. ACS Chem. Biol. 2016, 11 (12), 3461–3472.

74. George, D.; Friedman, M.; Allen, H.; Argiriadi, M.; Barberis, C.; Bischoff, A.; Clabbers, A.; Cusack, K.; Dixon, R.; Fix-Stenzel, S.; Gordon, T.; Janssen, B.; Jia, Y.; Moskey, M.; Quinn, C.;

75. Salmeron, J. A.; Wishart, N.; Woller, K.; Yu, Z. T., Discovery of thieno[2,3-c]pyridines as potent COT inhibitors. Bioorganic & Medicinal Chemistry Letters 2008, 18 (18), 4952–4955.

76. Koehler, M. F. T.; Bergeron, P.; Blackwood, E. M.; Bowman, K.; Clark, K. R.; Firestein, R.; Kiefer, J. R.; Maskos, K.; McCleland, M. L.; Orren, L.; Salphati, L.; Schmidt, S.; Schneider, E. V.; Wu, J. S.; Beresini, M. H., Development of a Potent, Specific CDK8 Kinase Inhibitor Which Phenocopies CDK8/19 Knockout Cells. Acs Medicinal Chemistry Letters 2016, 7 (3), 223–228.

77. Prosser, S. L.; Sahota, N. K.; Pelletier, L.; Morrison, C. G.; Fry, A. M., Nek5 promotes centrosome integrity in interphase and loss of centrosome cohesion in mitosis. J. Cell Biol. 2015, 209 (3), 339–348.

78. Prosser, S. L.; Fry, A. M., Nek5: a new regulator of centrosome integrity. Oncotarget 2015, 6 (28), 24594–5.

79. Melo Hanchuk, T. D.; Papa, P. F.; La Guardia, P. G.; Vercesi, A. E.; Kobarg, J., Nek5 interacts with mitochondrial proteins and interferes negatively in mitochondrial mediated cell death and respiration. Cell. Signalling 2015, 27 (6), 1168–1177.

80. Liu, Q.; Sabnis, Y.; Zhao, Z.; Zhang, T.; Buhrlage, S. J.; Jones, L. H.; Gray, N. S., Developing irreversible inhibitors of the protein kinase cysteinome. Chem Biol 2013, 20 (2), 146–59.

81. Yin, M.-J.; Shao, L.; Voehringer, D.; Smeal, T.; Jallal, B., The serine/threonine kinase Nek6 is required for cell cycle progression through mitosis. J. Biol. Chem. 2003, 278 (52), 52454–52460.

82. Rapley, J.; Nicolas, M.; Groen, A.; Regue, L.; Bertran, M. T.; Caelles, C.; Avruch, J.; Roig, J., The NIMA-family kinase Nek6 phosphorylates the kinesin Eg5 at a novel site necessary for mitotic spindle formation. J. Cell Sci. 2008, 121 (23), 3912–3921.

83. Lee, M.-Y.; Kim, H.-J.; Kim, M.-A.; Jee, H. J.; Kim, A. J.; Bae, Y.-S.; Park, J.-I.; Chung, J. H.; Yun, J., Nek6 is involved in G2/M phase cell cycle arrest through DNA damage-induced phosphorylation. Cell Cycle 2008, 7 (17), 2705–2709.

84. Takeno, A.; Takemasa, I.; Doki, Y.; Yamasaki, M.; Miyata, H.; Takiguchi, S.; Fujiwara, Y.; Matsubara, K.; Monden, M., Integrative approach for differentially overexpressed genes in gastric cancer by combining large-scale gene expression profiling and network analysis. Br. J. Cancer 2008, 99 (8), 1307–1315.

85. Jeon, Y. J.; Lee, K. Y.; Cho, Y.-Y.; Pugliese, A.; Kim, H. G.; Jeong, C.-H.; Bode, A. M.; Dong, Z., Role of NEK6 in tumor promoter-induced transformation in JB6 C141 mouse skin epidermal cells. J. Biol. Chem. 2010, 285 (36), 28126–28133.

86. Capra, M.; Nuciforo, P. G.; Confalonieri, S.; Quarto, M.; Bianchi, M.; Nebuloni, M.; Boldorini, R.; Pallotti, F.; Viale, G.; Gishizky, M. L.; Draetta, G. F.; Di Fiore, P. P., Frequent Alterations in the Expression of Serine/Threonine Kinases in Human Cancers. Cancer Research 2006, 66 (16), 8147.

87. Srinivasan, P.; Chella Perumal, P.; Sudha, A., Discovery of Novel Inhibitors for Nek6 Protein through Homology Model Assisted Structure Based Virtual Screening and Molecular Docking Approaches. The Scientific World Journal 2014, 2014, 9.

88. Moraes, E.; Meirelles, G.; Honorato, R.; de Souza, T.; de Souza, E.; Murakami, M.; de Oliveira, P.; Kobarg, J., Kinase Inhibitor Profile for Human Nek1, Nek6, and Nek7 and Analysis of the Structural Basis for Inhibitor Specificity. Molecules 2015, 20 (1), 1176.

89. Min, J.; Guo, K.; Suryadevara, P. K.; Zhu, F.; Holbrook, G.; Chen, Y.; Feau, C.; Young, B. M.; Lemoff, A.; Connelly, M. C.; Kastan, M. B.; Guy, R. K., Optimization of a Novel Series of Ataxia-Telangiectasia Mutated Kinase Inhibitors as Potential Radiosensitizing Agents. Journal of Medicinal Chemistry 2016, 59 (2), 559–577.

90. Beria, I.; Valsasina, B.; Brasca, M. G.; Ceccarelli, W.; Colombo, M.; Cribioli, S.; Fachin, G.; Ferguson, R. D.; Fiorentini, F.; Gianellini, L. M.; Giorgini, M. L.; Moll, J. K.; Posteri, H.; Pezzetta, D.; Roletto, F.; Sola, F.; Tesei, D.; Caruso, M., 4,5-Dihydro-1H-pyrazolo[4,3-h]quinazolines as potent and selective Polo-like kinase 1 (PLK1) inhibitors. Bioorganic & Medicinal Chemistry Letters 2010, 20 (22), 6489–6494.

91. Yissachar, N.; Salem, H.; Tennenbaum, T.; Motro, B., Nek7 kinase is enriched at the centrosome, and is required for proper spindle assembly and mitotic progression. FEBS Lett. 2006, 580 (27), 6489–6495.

92. Kim, S.; Lee, K.; Rhee, K., NEK7 is a centrosomal kinase critical for microtubule nucleation. Biochem. Biophys. Res. Commun. 2007, 360 (1), 56–62.

93. O'Regan, L.; Fry, A. M., The Nek6 and Nek7 protein kinases are required for robust mitotic spindle formation and cytokinesis. Mol. Cell. Biol. 2009, 29 (14), 3975–3990.

94. Belham, C.; Comb, M. J.; Avruch, J., Identification of the NIMA family kinases NEK6/7 as regulators of the p70 ribosomal S6 kinase. Curr. Biol. 2001, 11 (15), 1155–1167.

95. Salem, H.; Rachmin, I.; Yissachar, N.; Cohen, S.; Amiel, A.; Haffner, R.; Lavi, L.; Motro, B., Nek7 kinase targeting leads to early mortality, cytokinesis disturbance and polyploidy. Oncogene 2010, 29 (28), 4046–4057.

96. Schmid-Burgk, J. L.; Chauhan, D.; Schmidt, T.; Ebert, T. S.; Reinhardt, J.; Endl, E.; Hornung, V., A genome-wide CRISPR (clustered regularly interspaced short palindromic repeats) screen identifies NEK7 as an essential component of NLRP3 inflammasome activation. J. Biol. Chem. 2016, 291 (1), 103–109.

97. Xu, J.; Lu, L.; Li, L., NEK7: a novel promising therapy target for NLRP3-related inflammatory diseases. Acta Biochim Biophys Sin (Shanghai) 2016, 48 (10), 966–968.

98. He, Y.; Hara, H.; Nunez, G., Mechanism and Regulation of NLRP3 Inflammasome Activation. Trends Biochem. Sci. 2016, 41 (12), 1012–1021.

99. Bowers, A. J.; Boylan, J. F., Nek8, a NIMA family kinase member, is overexpressed in primary human breast tumors. Gene 2004, 328, 135–142.

100. Holland, P. M.; Milne, A.; Garka, K.; Johnson, R. S.; Willis, C.; Sims, J. E.; Rauch, C. T.; Bird, T. A.; Virca, G. D., Purification, cloning, and characterization of Nek8, a novel NIMA-related kinase, and its candidate substrate Bicd2. J. Biol. Chem. 2002, 277 (18), 16229–16240.

101. Abeyta, A.; Castella, M.; Jacquemont, C.; Taniguchi, T., NEK8 regulates DNA damage-induced RAD51 foci formation and replication fork protection. Cell Cycle 2017, 16 (4), 335–347.

102. Otto, E. A.; Trapp, M. L.; Schultheiss, U. T.; Helou, J.; Quarmby, L. M.; Hildebrandt, F., NEK8 mutations affect ciliary and centrosomal localization and may cause nephronophthisis. J. Am. Soc. Nephrol. 2008, 19 (3), 587–592.

103. Smith, L. A.; Bukanov, N. O.; Husson, H.; Russo, R. J.; Barry, T. C.; Taylor, A. L.; Beier, D. R.; Ibraghimov-Beskrovnaya, O., Development of polycystic kidney disease in juvenile cystic kidney mice: insights into pathogenesis, ciliary abnormalities, and common features with human disease. J. Am. Soc. Nephrol. 2006, 17 (10), 2821–2831.

104. Belham, C.; Roig, J.; Caldwell, J. A.; Aoyama, Y.; Kemp, B. E.; Comb, M.; Avruch, J., A Mitotic Cascade of NIMA Family Kinases: Nercc1/Nek9 Activates THE Nek6 AND Nek7 Kinases. J. Biol. Chem. 2003, 278 (37), 34897–34909.

105. Tan, B. C.-M.; Lee, S.-C., Nek9, a novel FACT-associated protein, modulates interphase progression. J. Biol. Chem. 2004, 279 (10), 9321–9330.

106. Yang, S.-W.; Gao, C.; Chen, L.; Song, Y.-L.; Zhu, J.-L.; Qi, S.-T.; Jiang, Z.-Z.; Wang, Z.-W.; Lin, F.; Huang, H.; Xing, F.-Q.; Sun, Q.-Y., Nek9 regulates spindle organization and cell cycle progression during mouse oocyte meiosis and its location in early embryo mitosis. Cell Cycle 2012, 11 (23), 4366–4377.

107. Kaneta, Y.; Ullrich, A., NEK9 depletion induces catastrophic mitosis by impairment of mitotic checkpoint control and spindle dynamics. Biochem. Biophys. Res. Commun. 2013, 442 (3-4), 139–146.

108. Levinsohn, J. L.; Sugarman, J. L.; McNiff, J. M.; Antaya, R. J.; Choate, K. A., Somatic Mutations in NEK9 Cause Nevus Comedonicus. Am. J. Hum. Genet. 2016, 98 (5), 1030–1037.

109. Cheung, M.; Kuntz, K. W.; Pobanz, M.; Salovich, J. M.; Wilson, B. J.; Andrews Iii, C. W.; Shewchuk, L. M.; Epperly, A. H.; Hassler, D. F.; Leesnitzer, M. A.; Smith, J. L.; Smith, G. K.; Lansing, T. J.; Mook Jr, R. A., Imidazo[5,1-f][1,2,4]triazin-2-amines as novel inhibitors of polo-like kinase 1. Bioorganic & Medicinal Chemistry Letters 2008, 18 (23), 6214–6217.

110. Moniz, L. S.; Stambolic, V., Nek10 mediates G2/M cell cycle arrest and MEK autoactivation in response to UV irradiation. Mol Cell Biol 2011, 31 (1), 30–42.

111. Greenman, C.; Stephens, P.; Smith, R.; Dalgliesh, G. L.; Hunter, C.; Bignell, G.; Davies, H.; Teague, J.; Butler, A.; Stevens, C.; Edkins, S.; O'Meara, S.; Vastrik, I.; Schmidt, E. E.; Avis, T.; Barthorpe, S.; Bhamra, G.; Buck, G.; Choudhury, B.; Clements, J.; Cole, J.; Dicks, E.; Forbes, S.; Gray, K.; Halliday, K.; Harrison, R.; Hills, K.; Hinton, J.; Jenkinson, A.; Jones, D.; Menzies, A.; Mironenko, T.; Perry, J.; Raine, K.; Richardson, D.; Shepherd, R.; Small, A.; Tofts, C.; Varian, J.; Webb, T.; West, S.; Widaa, S.; Yates, A.; Cahill, D. P.; Louis, D. N.; Goldstraw, P.; Nicholson, A. G.; Brasseur, F.; Looijenga, L.; Weber, B. L.; Chiew, Y. E.; DeFazio, A.; Greaves, M. F.; Green, A. R.; Campbell, P.; Birney, E.; Easton, D. F.; Chenevix-Trench, G.; Tan, M. H.; Khoo, S. K.; Teh, B. T.; Yuen, S. T.; Leung, S. Y.; Wooster, R.; Futreal, P. A.; Stratton, M. R., Patterns of somatic mutation in human cancer genomes. Nature 2007, 446 (7132), 153–8.

112. Smith, C. R.; Dougan, D. R.; Komandla, M.; Kanouni, T.; Knight, B.; Lawson, J. D.; Sabat, M.; Taylor, E. R.; Vu, P.; Wyrick, C., Fragment-Based Discovery of a Small Molecule Inhibitor of Bruton's Tyrosine Kinase. Journal of Medicinal Chemistry 2015, 58 (14), 5437–5444.

113. Smith, D. G.; Buffet, M.; Fenwick, A. E.; Haigh, D.; Ife, R. J.; Saunders, M.; Slingsby, B. P.; Stacey, R.; Ward, R. W., 3-Anilino-4-arylmaleimides: potent and selective inhibitors of glycogen synthase kinase-3 (GSK-3). Bioorganic & Medicinal Chemistry Letters 2001, 11 (5), 635–639.

114. Noguchi, K.; Fukazawa, H.; Murakami, Y.; Uehara, Y., Nucleolar Nek11 Is a Novel Target of Nek2A in G1/S-arrested Cells. J. Biol. Chem. 2004, 279 (31), 32716–32727.

115. Melixetian, M.; Klein, D. K.; Sorensen, C. S.; Helin, K., NEK11 regulates CDC25A degradation and the IR-induced G2/M checkpoint. Nat. Cell Biol. 2009, 11 (10), 1247–1253.

116. Sabir, S. R.; Sahota, N. K.; Fry, A. M.; Jones, G. D. D., Loss of Nek11 Prevents G2/M Arrest and Promotes Cell Death in HCT116 Colorectal Cancer Cells Exposed to Therapeutic DNA Damaging Agents. PLoS One 2015, 10 (10), e0140975.

117. Piao, S.; Lee, S.-J.; Xu, Y.; Gwak, J.; Oh, S.; Park, B.-J.; Ha, N.-C., CK1ε targets Cdc25A for ubiquitin-mediated proteolysis under normal conditions and in response to checkpoint activation. Cell Cycle 2011, 10 (3), 531–537.

118. Soerensen, C. S.; Melixetian, M.; Klein, D. K.; Helin, K., NEK11: Linking CHK1 and CDC25A in DNA damage checkpoint signaling. Cell Cycle 2010, 9 (3), 450–455.

119. Noguchi, K.; Fukazawa, H.; Murakami, Y.; Uehara, Y., Nek11, a New Member of the NIMA Family of Kinases, Involved in DNA Replication and Genotoxic Stress Responses. J. Biol. Chem. 2002, 277 (42), 39655–39665.

120. Tsai, J.; Lee, J. T.; Wang, W.; Zhang, J.; Cho, H.; Mamo, S.; Bremer, R.; Gillette, S.; Kong, J.; Haass, N. K.; Sproesser, K.; Li, L.; Smalley, K. S. M.; Fong, D.; Zhu, Y.-L.; Marimuthu, A.; Nguyen, H.; Lam, B.; Liu, J.; Cheung, I.; Rice, J.; Suzuki, Y.; Luu, C.; Settachatgul, C.; Shellooe, R.; Cantwell, J.; Kim, S.-H.; Schlessinger, J.; Zhang, K. Y. J.; West, B. L.; Powell, B.; Habets, G.; Zhang, C.; Ibrahim, P. N.; Hirth, P.; Artis, D. R.; Herlyn, M.; Bollag, G., Discovery of a selective inhibitor of oncogenic B-Raf kinase with potent antimelanoma activity. Proceedings of the National Academy of Sciences 2008, 105 (8), 3041–3046.

121. Bollag, G.; Hirth, P.; Tsai, J.; Zhang, J.; Ibrahim, P. N.; Cho, H.; Spevak, W.; Zhang, C.; Zhang, Y.; Habets, G.; Burton, E. A.; Wong, B.; Tsang, G.; West, B. L.; Powell, B.; Shellooe, R.; Marimuthu, A.; Nguyen, H.; Zhang, K. Y. J.; Artis, D. R.; Schlessinger, J.; Su, F.; Higgins, B.; Iyer, R.; D/'Andrea, K.; Koehler, A.; Stumm, M.; Lin, P. S.; Lee, R. J.; Grippo, J.; Puzanov, I.; Kim, K. B.; Ribas, A.; McArthur, G. A.; Sosman, J. A.; Chapman, P. B.; Flaherty, K. T.; Xu, X.; Nathanson, K. L.; Nolop, K.,

122. Clinical efficacy of a RAF inhibitor needs broad target blockade in BRAF-mutant melanoma. Nature 2010, 467 (7315), 596–599.

123. Rheault, T. R.; Stellwagen, J. C.; Adjabeng, G. M.; Hornberger, K. R.; Petrov, K. G.; Waterson, A. G.; Dickerson, S. H.; Mook, R. A.; Laquerre, S. G.; King, A. J.; Rossanese, O. W.; Arnone, M. R.; Smitheman, K. N.; Kane-Carson, L. S.; Han, C.; Moorthy, G. S.; Moss, K. G.; Uehling, D. E., Discovery of Dabrafenib: A Selective Inhibitor of Raf Kinases with Antitumor Activity against B-Raf-Driven Tumors. ACS Medicinal Chemistry Letters 2013, 4 (3), 358–362.

